# Distinct Roles of Kif6 and Kif9 in Mammalian Ciliary Trafficking and Motility

**DOI:** 10.1101/2023.11.09.564704

**Authors:** Chuyu Fang, Xinwen Pan, Di Li, Yawen Chen, Luan Li, Qi Gao, Dong Li, Xueliang Zhu, Xiumin Yan

## Abstract

Ciliary beat and intraflagellar transport (IFT) depend on dynein and kinesin motors. Kinesin-9 family members Kif6 and Kif9 are implicated in ciliary motilities across protists and mammals. How they function and whether they act redundantly, however, remain unclear. Here, we show that they play distinct roles in mammals. Kif6 forms puncta that move bidirectionally without or with IFT-B particles along axonemes, whereas Kif9 is immobilized on ciliary central apparatus. Only Kif6 binds to and glides microtubules, and the activities are self-inhibited. *Kif6* deficiency in mice impairs directional ciliary beat across ependymal tissues and cerebrospinal fluid flow, resulting in severe hydrocephalus and high mortality, whereas *Kif9* deficiency induces mild hydrocephalus without obviously defective ciliary beat and life span. Both *Kif6^-/-^* and *Kif9^-/-^* males are infertile but show respectively oligozoospermia with poor sperm motility and defective forward motion of sperms. These results suggest Kif6 as a motile cilia-specific IFT motor and Kif9 as a central apparatus regulator.

## Introduction

Cilia are evolutionarily conserved, microtubule (MT)-based organelles that protrude from the cell surface and are generally categorized into primary and motile cilia (Klena and Pigino, 2022; Reiter and Leroux, 2017). Primary cilia are usually possess 9+0 MT arrangement consisting of nine peripheral doublet MTs (DMTs), whereas most motile cilia/flagella typically possess a 9+2 MT axoneme with a central pair (CP) of MT singlets (C1 and C2) surrounded by nine DMTs (Han et al., 2022; Ishikawa, 2017; Loreng and Smith, 2017). The CP MTs are decorated with proteinaceous projections and the peripheral DMTs are equipped with dynein arms and radial spokes. The signal from CP apparatus transmits through radial spokes to govern the sliding of axonemal dyneins along adjacent MTs, generating cilia/flagella beating (Gui et al., 2021; Hoek et al., 2022; Loreng and Smith, 2017). Except for sperm flagella, motile cilia are mainly present as multicilia with dozens to hundreds of cilia within a single cell in mammals. Multicilia are found in epithelia of various organs such as the brain ventricles, the airway and the oviduct, playing critical roles in cerebrospinal fluid (CSF) circulation, mucus clearance, and ovum transport (Legendre et al., 2021; Marshall and Kintner, 2008; Tissir et al., 2010).

To generate directional fluid flow, proper polarities are needed to establish at both cellular and tissue levels. During the differentiation of multiciliated cells, basal bodies (BBs) initially amplify and dock to the cell membrane, as indicated by the orientiation of their basal feet (BFs), which are conical projections on BB walls that align with the direction of effective strokes (Gibbons, 1961; Mirzadeh et al., 2010; Ohata and Alvarez-Buylla, 2016). As maturation progress, the orientation of BBs within individual cells gradually aligns in a coordinated manner along the direction of fluid flow, establishing rotational polarity (Boutin et al., 2014; Guirao et al., 2010). In ependymal cells, in addition to rotational polarity, BBs exhibit a specific clustering into a patch on one side of the cell surface, establishing translational polarity that distinguishes them from other multiciliated cells such as tracheal epithelial cells (Mirzadeh et al., 2010). These processes ultimately establish the planar polarity of multiciliated tissues. The establishment of planar polarity involves planar cell polarity (PCP) pathway and hydrodynamic force flow, guiding the process over several days (Boutin et al., 2014; Ohata and Alvarez-Buylla, 2016).

Kinesins are molecular motors that play essential roles in directional transport of cargos through processive movements along MTs or regulation of MT dynamics in various cellular processes such as mitosis, ciliogenesis and intracellular transport across eukaryotes (Cason and Holzbaur, 2022; Hirokawa et al., 2009; Lu and Gelfand, 2017). They are classified into N-kinesins, M-kinesins, and C-kinesins based on the position of motor domain. Two kinesin families, kinesin-2 and 9, have been found to function in cilia or flagella. The kinesin-2 family motors (Kif3a/b/c) are well-known for their involvement in IFT, a specialized bidirectional transportation machinery (Klena and Pigino, 2022; Scholey, 2013). IFT relies on two multi-subunit complexes, IFT-A and IFT-B, which are powered by kinesin-2 or cytoplasmic dynein-2 respectively for anterograde or retrograde trafficking along axonemal MTs in the form of IFT trains (Nakayama and Katoh, 2018). IFT enables the transport of ciliary proteins in and out of cilia, critical for cilia assembly, maintenance, and function (Meleppattu et al., 2022; Nakayama and Katoh, 2018; Pigino, 2021).

The kinesin-9 family consists of two members, Kif6 and Kif9, which share a high sequence identity at their motor domains. They belong to the N-kinesin group, typically involved in plus-end-directed transports of cargos (Hirokawa et al., 2009; Lu and Gelfand, 2017). Unlike the kinesin-2 family, which plays a critical role in both primary and motile cilia, the kinesin-9 family is predominantly associated with flagellated species (Demonchy et al., 2009; Scholey, 2013), suggesting distinct or redundant roles in regulating flagella motility. In *Chlamydomonas*, only KLP1, the homolog of Kif9, has been studied. It specifically localizes to the CP MTs and is proposed as an active motor to regulate flagellar motility (Bernstein et al., 1994; Han et al., 2022; Yokoyama et al., 2004). Interestingly, in *Trypanosoma*, the kinesin-9 family has been demonstrated distinct functions in flagella motility (Demonchy et al., 2009). Knockdown of *KIF9A*, the *Trypanosoma* homolog of *Kif9*, reduces flagellum beating and cell movement, while knockdown of *KIF9B*, the homolog of *Kif6*, results in cell paralysis and defective assembly of the paraflagellar rod (PFR). Although there is no observed movement of KIF9B within flagella, it is speculated that it is involved in PFR transport or the recognition of axoneme asymmetry for PFR assembly in *Trypanosoma* (Demonchy et al., 2009). In mammals, mutations in *Kif9* have been linked to defective sperm motility and impaired fertility (Meng et al., 2022; Miyata et al., 2020). However, its function in motile cilia remains unexplored. A frameshift mutation of *KIF6* has been found in a patient with intellectual disability and megalocephaly (Konjikusic et al., 2018). Mice carrying a *Kif6* C-terminal truncation mutant (*Kif6^p.G555fs^*) develop severe hydrocephalus possibly due to defective ciliogenesis in the ependyma (Konjikusic et al., 2018). Since the mutant mice still express the motor domain of Kif6, it cannot be ruled out that the observed hydrocephalus phenotype and defective ciliogenesis could be caused by the dominant negative effect of the remaining motor domain. Despite these achievements, a comprehensive understanding of the roles of Kif6 and Kif9 in mammalian ciliary trafficking and cilia motility is still lacking.

In this study, we investigate the functions of Kif6 and Kif9 in mice. Our results suggest that Kif6 is a motile cilia-specific motor involved in IFT and essential for the planar polarity of multicilia in the ependyma, whereas Kif9 functions in the central apparatus for regulating cilia/flagella motility.

## Results

### Mammalian Kif6 and Kif9 are motile cilia-specific kinesins with distinct localizations

Murine full-length Kif6 (NP_796026.2) and Kif9 (NP_001157041.1) are highly conserved proteins composed of 802 and 810 amino acid residues, respectively. Structurally, both proteins have a motor domain located at the N-terminus, followed by two coiled-coil regions and a tail at the C-terminus (Fig. 1a). Their motor domains share a sequence identity of 43%, while their C-termini are less conserved (Fig.1a and Supplementary Fig. 1a, b). Phylogenetic analysis showed that Kif6 and Kif9 co-existed in eukaryotes with motile cilia/flagella, dating back to protists like *Chlamydomonas* and *Tetrahymena* (Fig. 1b and Supplementary Fig. 1a) (Bernstein et al., 1994; Vincensini et al., 2011; Yokoyama et al., 2004). In contrast, they lacked homologues in *Caenorhabditis elegans*, which only contains immotile sensory cilia (Fig. 1b) (Inglis et al., 2007; Vincensini et al., 2011). These data imply that Kif6 and Kif9 are evolved to be both important for motile cilia.

**Figure 1.**
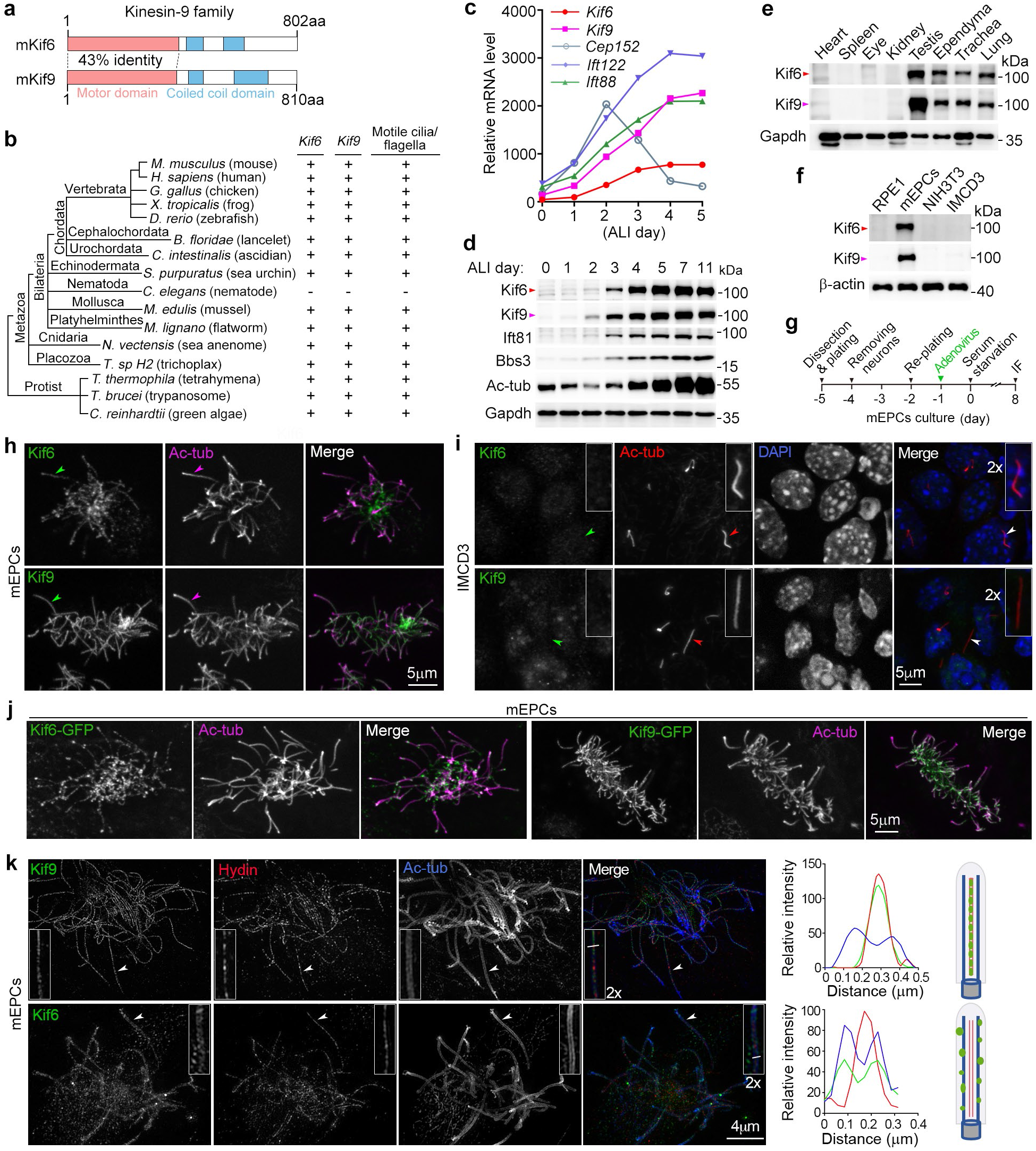
Molecular characterizations and subcellular localizations of mammalian *Kif6* and *Kif9*. **(a)** Schematics of full-length murine Kif6 and Kif9. The motor and coiled-coil domains were identified using Conserved Domain Search Service of NCBI and SMART (Simple Modular Architecture Research Tool) (http://smart.embl.de). **(b)** Conservation of *Kif6* and *Kif9* across species with motile cilia and/or flagella throughout the evolution process. The phylogenetic tree and taxonomic groups were referrenced from literature (Cetkovic et al., 2018; Mukherjee and Brocchieri, 2013). **(c)** mRNA expression profiles of *Kif6* and *Kif9* during multiciliation. The gene expression profiles were obtained from our previous cDNA microarray analysis of mTECs cultured at an air-liquid interface (ALI) for the indicated days (Xu et al., 2015). Expression patterns of genes crucial for cilia formation (*Ift88* and *Ift122*) and BB amplification (*Cep152*), were listed for comparison (Klena and Pigino, 2022; Zhao et al., 2013). **(d)** Expression patterns of Kif6 and Kif9 during multiciliation of mTECs. Multicilia formation is indicated by increased levels of Ift81, Bbs3 and acetylated tubulin (Ac-tub) (Klena and Pigino, 2022). Gapdh served as loading control. **(e, f)** Abundance of Kif6 and Kif9 in mouse tissues (**e**) and cultured cells (**f**) containing motile cilia. Tissues from a 2-month-old male mouse were dissected and lysed. mEPCs were harvested on day 8 post serum starvation. RPE1, NIH3T3 and IMCD3 cells were serum-starved for 48 h to induce ciliogenesis. Gapdh or β-actin served as loading control. **(g)** Experimental scheme for **h, j, k** and Supplementary Fig. 1c. Precursors of mEPCs were isolated from P0 mouse brain and serum-starved on day 0 to induce to differentiate into multiciliated cells. To express GFP-tagged proteins, the cells were infected with adenovirus on day-1. The cells were fixed on day 8 for immunofluorescence (IF) staining and imaged by confocal microscopy (**h**, **j**) or 3D-SIM (**k**). **(h)** Localizations of endogenous Kif6 and Kif9 in motile cilia. Ac-tub labeled ciliary axonemes. Arrowheads indicate typical cilia. **(i)** Kif6 and Kif9 did not localize in primary cilia. IMCD3 cells were serum-starved to induce primary ciliogenesis and fixed at day 2 post serum starvation for IF, followed by confocal microscopy. The nucleus was visualized by DAPI, a DNA-specific dye. Cilia pointed by arrowheads were zoomed in 2× to show details. **(j)** Exogenous Kif6-GFP and Kif9-GFP localized at the axonemes of multicilia. **(k)** Typical 3D-SIM images of endogenous Kif9 and Kif6. Ac-tub and Hydin served as axoneme and CP markers, respectively. Cilia pointed were magnified by 200% for details. Corresponding line scans at the indicated region of the magnified insets showed colocalizations of Kif9 with Hydin and of Kif6 with Ac-tub. Illustrations are provided to aid comprehensions.

We re-analyzed our previous microarray results (Xu et al., 2015) and found that *Kif6* and *Kif9* were highly upregulated during the multiciliation of mouse tracheal epithelial cells (mTECs) (Fig. 1c). Immunoblots confirmed a substantial increase in the protein levels of Kif6 and Kif9 during multiciliation of mTECs (Fig. 1d). Furthermore, Kif6 and Kif9 were specifically expressed in tissues and cells abundant in motile cilia or flagella, such as the ependyma, the trachea, the lung, the testis, and primary cultured mouse ependymal cells (mEPCs), but were hardly detected in tissues and cells with only immotile cillia (Fig. 1e, f). Consistently, immunostaining indicated their localizations in motile cilia of mEPCs (Fig. 1g, h) but not in primary cilia of IMCD3 cells (Fig. 1i). To validate their ciliary localization, we infected mEPCs with adenovirus to express Kif6-GFP or Kif9-GFP (Fig. 1g) and clearly observed localizations of the exogenous proteins in motile cilia (Fig. 1j).

To visualize their detailed localizations, we performed three dimensional structured illumination microscopy (3D-SIM). Ciliary Kif9 localized in the central lumen of axonemes marked with acetylated tubulin (Ac-tub) and colocalized well with the CP marker Hydin (Fig. 1k) (Lechtreck et al., 2008), suggesting a conserved localization in the C2 projection as *Chlamydomonas* KLP1 (Bernstein et al., 1994; Han et al., 2022; Yokoyama et al., 2004). In sharp contrast, Kif6 mainly localized along the axonemes as puncta (Fig. 1k). Similar ciliary localization patterns were observed for GFP-tagged Kif6 and Kif9, using Spef1 as a CP marker (Supplementary Fig. 1c) (Zheng et al., 2019). Therefore, Kif6 and Kif9 appear to play different roles speficifically in motile cilia.

### Kif6, but not Kif9, displays IFT-like behaviors along ciliary axonemes

As kinesins with a motor domain at their N-terminus typically function as plus-end-directed motor proteins (Cason and Holzbaur, 2022; Hirokawa et al., 2009), we sought to examine whether GFP-tagged Kif6 and Kif9 expressed in mEPCs (Fig. 2a) could move along DMTs and central MTs, respectively. Immunoblotting demonstrated comparable expression levels of GFP-tagged Kif6 and Ift81, an IFT-B component (Klena and Pigino, 2022), with their endogenous proteins, whereas Kif9-GFP showed relatively higher expression than endogenous Kif9 (Fig. 2b). To achieve superior spatiotemporal resolution, we employed grazing incidence structured illumination microscopy (GI-SIM) (Guo et al., 2018; Qiao et al., 2022). As shown previously (Qiao et al., 2022), Ift81-GFP particles displayed clear bidirectional movements along axonemes (Fig. 2c and Supplementary Fig. 2a; Movie 1). Strikingly, Kif6-GFP puncta also underwent robust and processive bidirectional movements resembling the IFT (Fig. 2d and Supplementary Fig. 2a; Movie 2). Kif9-GFP, however, did not exhibit any processive movement along axonemes (150 cilia from 10 cells) (Fig. 2e and Supplementary Fig. 2a; Movie 3), suggesting that it is immobilized on the central apparatus.

**Figure 2.**
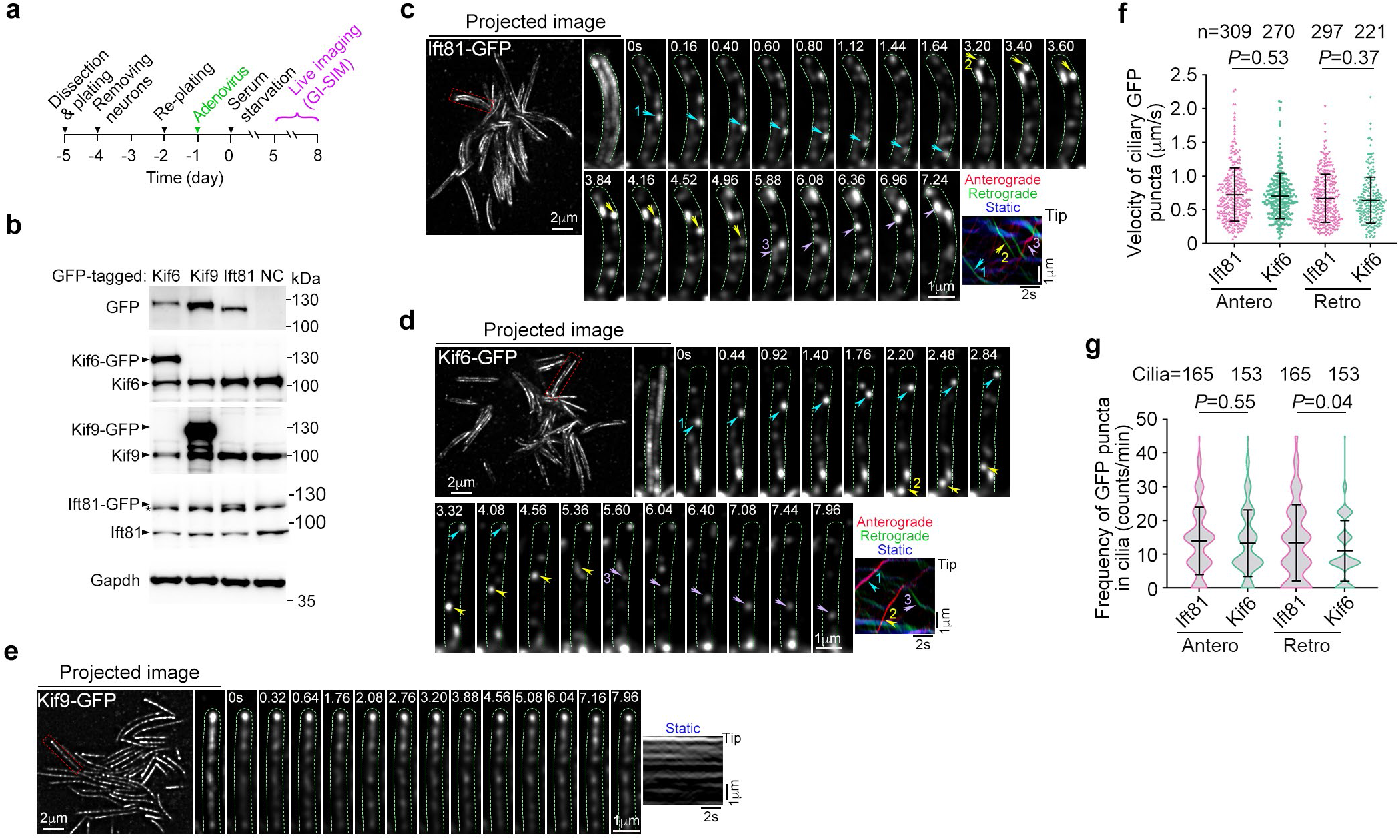
Kif6 moves along axonemes in IFT-like pattern and Kif9 is immobilized on the central apparatus. Quantification results are presented as mean ± SD and subjected to unpaired two-tailed student’s *t*-test. **(a)** Experimental scheme for ciliary trafficking recorded by GI-SIM. mEPCs were infected with adenovirus at day-1 and imaged during days 5-8. **(b)** Expression levels of GFP-tagged proteins in day 7 mEPCs. Immunoblotting was performed using GFP, Kif6, Kif9 and Ift81 antibodies, respectively. A star indicates a non-specific band closed to Ift81-GFP. **(c-e)** Ciliary movement of Ift81-GFP (**c**), Kif6-GFP (**d**) and Kif9-GFP (**e**). Both Ift81-GFP (**c**) and Kif6-GFP (**d**) were able to move along the axonemal outer DMTs while the motility of Kif9-GFP (**e**) was not detectable. Images were recorded at 25 frames per sec (fps). The trafficking trajectories of GFP-tagged puncta through time projections of the first 200 frames from Movies 1-3 were showed. Typical cilia framed by dashed red lines in projected images, were magnified by 300% to show details. Dash green lines were used to outline the cilia. The clear and traceable particles were marked respectively by arrowheads for anterograde and by arrows for retrograde to show the progressive movements. The corresponding kymographs were presented. (**f, g**) Quantification results of velocities and frequencies from the indicated numbers of particles (**f**) and cilia (**g**). Antero: Anterograde transport; Retro: Retrograde transport; s: second.

Qunatification results revealed that the Kif6-GFP puncta moved at 0.71 ± 0.34 μm/s anterogradely and 0.64 ± 0.34 μm/s retrogradely, similar to those of the Ift81-GFP particles (0.73 ± 0.40 μm/s anterogradely and 0.67 ± 0.36 μm/s retrogradely) (Fig. 2f). Frequencies of movements were also similar between the two (13.3 ± 11.3 and 14.0 ± 10.0 counts/min respectively for anterograde Kif6-GFP puncta and Ift81-GFP particles; 11.0 ± 9.0 and 13.2 ± 9.9 counts/min for retrograde Kif6-GFP puncta and Ift81-GFP particles) (Fig. 2g). These results suggest that Kif6 is involved in IFT trafficking.

### Ciliary Kif6 puncta traffic with or without IFT-B particles

To understand the relationship between Kif6 and IFT, we immunostained Kif6-GFP-expressing mEPCs to visualize IFT-B component Ift88 or Ift56 (Klena and Pigino, 2022). 3D-SIM images showed that some Kif6 puncta were located adjacent to IFT-B particles while the others appeared to be alone (Fig. 3a and Supplementary Fig. 2b).

**Figure 3.**
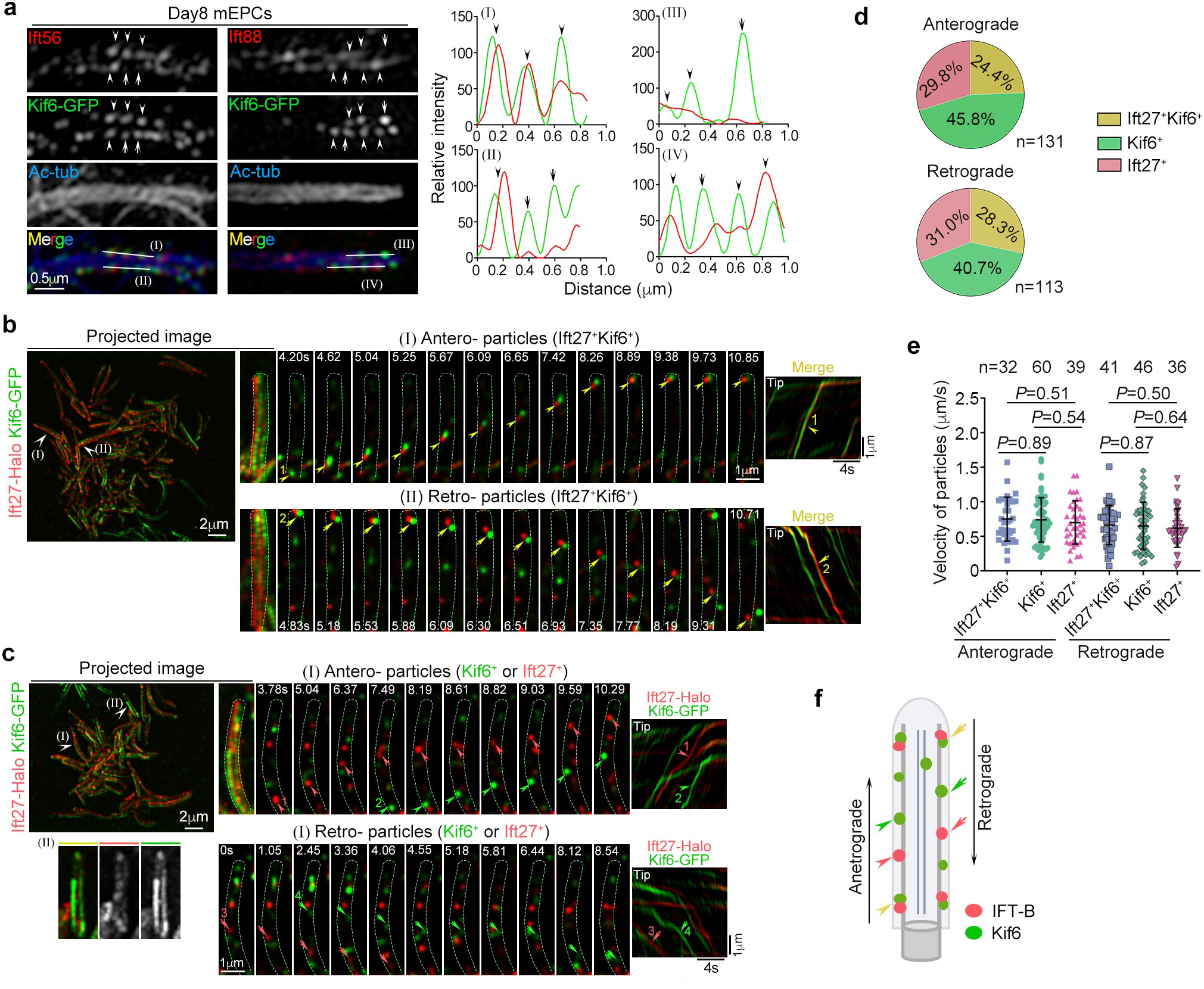
Kif6-GFP puncta move with or without IFT-B particles along the DMTs. **(a)** Typical 3D-SIM images of Kif6-GFP and IFT proteins (Ift56 and Ift88). Cilia cropped from Supplementary Fig. 2b were magnified by 400% to show details. Ac-tub marked ciliary axoneme. Line scans at the indicated regions (I-IV) exihibited the Kif6-GFP with or without IFT proteins, which were respectively pointed by arrowheads or arrows. **(b-f)** Ciliary trafficking of fluorescent particles. mEPCs were infected with adenovirus expressing of Kif6-GFP and Ift27-Halo and were performed dual-color time-lapse live imaging at 14 fps. Three distinct bidirectional moving particles within cilia were observed: labeled with both Ift27-Halo and Kif6-GFP (Ift27^+^Kif6^+^ duo-particles), only with Kif6-GFP (Kif6^+^ particles) and only with Ift27-Halo (Ift27^+^ particles). Ciliary trafficking trajectories of GFP/Halo-tagged puncta were projected by the first 200 frames from Movies 4-5. Typical trajectories pointed were magnified by 300% to roughly outline the cilia in projected images. The clear and traceable particles were indicated to display anterograde (arrowheads) and retrograde (arrows) transports of Ift27^+^Kif6^+^ duo-particles (**b**), Kif6^+^ particles (**c**) and Ift27^+^ particles (**c**) from representative time sequences. Dashed green lines were used to outline the cilia. Corresponding kymographs were presented. The ratios (**d**) and velocities (**e**) of these three distinct trafficking particles were quantified from 45 cilia in 4 mEPCs. The number (n) of corresponding particles was presented. Refer to Supplementary Fig. 2d for additional example of Kif6^+^ or Ift27^+^ bidirectional movements. Illustration is provided to aid comprehensions (**f**).

Next, we co-expressed Kif6-GFP and Ift27-Halo, another IFT-B component (Klena and Pigino, 2022), in mEPCs by adenoviral infection and performed dual-color time-lapse GI-SIM imaging. To ensure proper alignment of different fluorescent channels, the microscopic system was calibrated routinely with fluorescent beads prior to imaging (Supplementary Fig. 2c). Interestingly, we observed three distinct types of bidirectional movements within cilia: co-moved Kif6-GFP puncta and Ift27-Halo particles (Ift27^+^ Kif6^+^ duo-particles), Kif6-GFP puncta alone (Kif6^+^ particles), and Ift27-Halo particles alone (Ift27^+^ particles) (Fig. 3b-d and Supplementary Fig. 2d; Movies 4-5). The moving Kif6^+^ puncta usually left two separate tracks in cilia, similar to the moving Ift27^+^ particles (Fig. 3b, c and Supplementary Fig. 2d), indicating that their movements are along peripheral doublet MTs instead of central MTs.

Quantification analyses on 45 cilia in 4 mEPCs indicated that ratios of these three trafficking types were approximately 1:1.5:1 for both anterograde and retrograde movements (Fig. 3d). These three types of particles exhibited comparable velocities: 0.75 ± 0.32 μm/s (Ift27^+^ Kif6^+^), 0.74 ± 0.32 μm/s (Kif6^+^), and 0.70 ± 0.31 μm/s (Ift27^+^) anterogradely, or 0.66 ± 0.28 μm/s, 0.65 ± 0.34 μm/s, and 0.62 ± 0.28 μm/s retrogradely (Fig. 3e), similar to those measured from mEPCs expressing only one exogenous protein (Fig. 2f).

These results suggest that Kif6 traffics bidrectionally in motile cilia as a discrete punctum either with or without an IFT-B particle (Fig. 3f).

### Kif6 is a self-inhibited MT-based motor

The trafficking beheviors of Kif6 (Fig. 3) suggested two possibilities: (1) Kif6 might function actively as a processive motor to transport cargos in motile cilia; and (2) Kif6 might only be transported as a passive cargo by IFT trains. To discriminate between these possibilities, we investigated its motor properties. Processive kinesins are self-inhibited by their C-termini and activated either physiologically through cargo binding or artifically by the removal of inhibitory domains (Cason and Holzbaur, 2022; Hirokawa et al., 2009). We thus transiently expressed GFP-Flag-tagged Kif6 or different truncation constructs containing the motor domain in HEK293T cells (Fig. 4a, b) to examine whether Kif6 could bind to MTs *in vivo* and whether the binding was regulated in a self-inhibitory manner. We observed that all the truncated constructs (Kif6-CC1, Kif6-CC2, and Kif6-MD) exhibited strong MT localizations, especially on the spindle of mitotic cells (Fig. 4c). Full-length Kif6, however, did not show MT localizations (Fig. 4c). In contrast, neither Kif9 nor its truncation constructs (Kif9-CC1, Kif9-CC2, and Kif9-MD) displayed MT associations (Fig. 4a, c and Supplementary Fig. 3a, b). To clarify whether the robust MT association *in vivo* was a direct effect, we purified the proteins from HEK293T cells (Fig. 3d) and performed MT binding assays *in vitro* (Fig. 4e) (Diao et al., 2022). Consistently, Kif6-CC1, Kif6-CC2, Kif6-MD, and Kif5c-MD, a positive control (Diao et al., 2022), all associated with MTs immobilized on glass coverslips, whereas full-length Kif6 and Kif9-CC1 still did not bind to the MTs (Fig. 4f). We next examined whether Kif6-CC1 could actively move along MTs through MT gliding assays (Fig. 4g) (Stanhope and Ross, 2015) and observed that, similar to Kif5c-MD, Kif6-CC1 tethered on coverslips also drove MT gliding (Fig. 4h and Movie 6). Therefore, Kif6 contains an MT-based motor activity that is autoinhibited by its C-terminus and activated presumbly by cargo-binding, supporting its involvement in axonemal MT-based cargo transport in cilia. On the other hand, Kif9’s lack of detectable MT-binding or motor activities (Fig. 4) also complies with its immotility on the central apparatus (Fig. 2e), further strengthening its role as an immobilized CP component.

**Figure 4.**
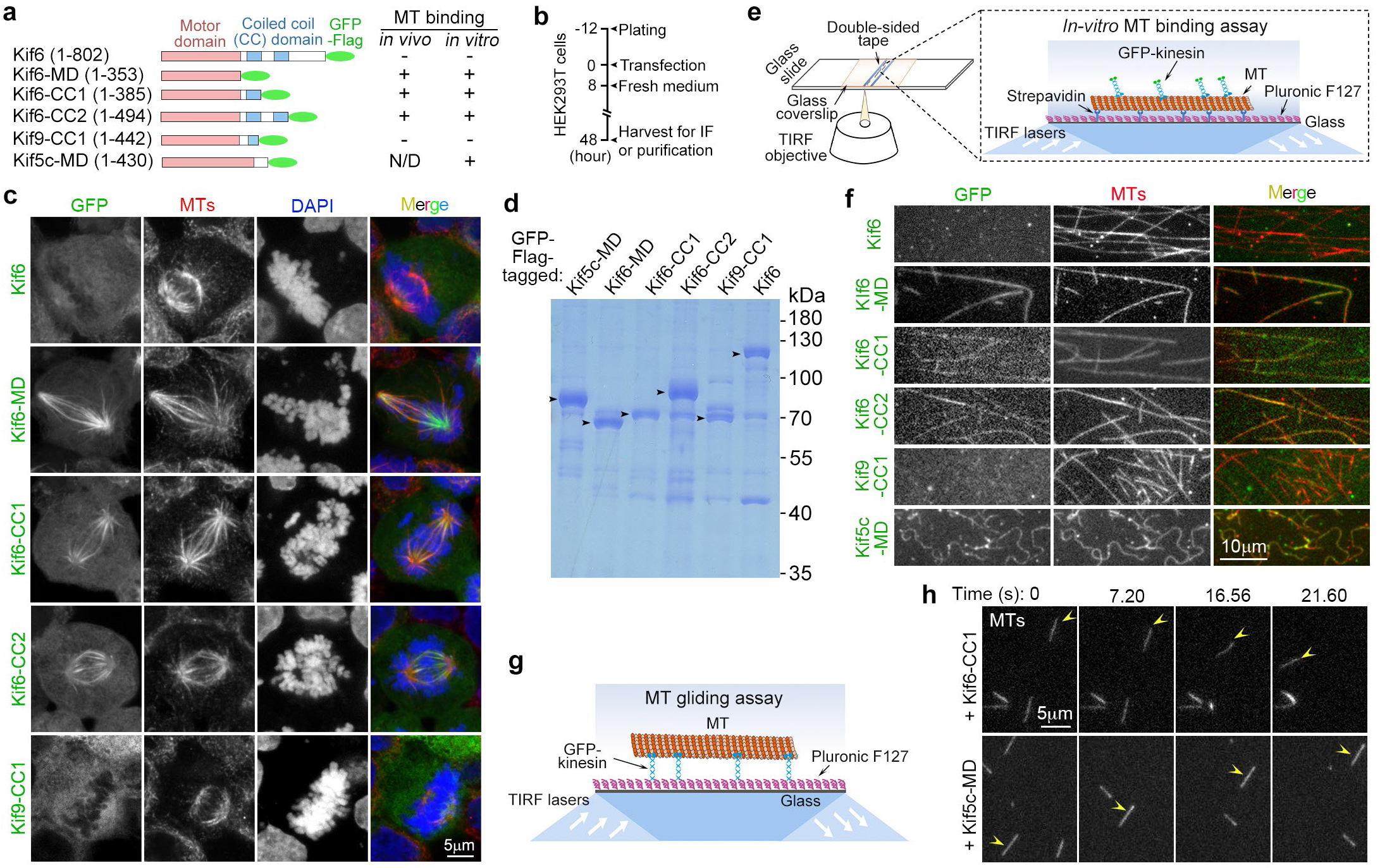
Kif6 but not Kif9 binds and glides MTs. **(a)** Diagrams of Kif6 and its deletion constructs showing their MT-binding ability. Kif5c-MD (1-430 aa) served as a positive control (Diao et al., 2022; Rogers et al., 2001). N/D, not determine. **(b)** Schematic of HEK293T cells culture and following analyses. HEK293T cells were transfected for 48 h and harvested for immunostaining (**c**) or protein purification (**d**). **(c)** Motor domain of Kif6 was required for spindle MTs binding. MTs were visualized by an anti-α-tubulin antibody. Chromosomes and nuclei were stained with DAPI. **(d)** Commassie blue-staining of purified proteins. Kif5c-MD was purified to serve as a positive control. **(e)** Schematic of *in-vitro* MT binding assays using total internal reflection fluorescence (TIRF) microscopy. Fluorescently labeled GMPCPP-stabilized microtubules immobilized on a Pluronic F127-treated glass coverslip via strepavidin linkages were imaged after the addition of GFP-tagged kinesin (final concentration: 0.5 μM) and ATP (1 mM). **(f)** The motor domain of Kif6, but not Kif9, bounds to MTs *in vitro*. **(g)** Schematic of *in-vitro* MT gliding assays using TIRF microscopy. Kif6-CC1 or Kif5c-MD were pre-immobilized on a coverslip and fluorescently labeled MTs were added for the assay. **(h)** Kif6-CC1 drove MT gliding. Arrowheads pointed to the moving MTs. Representative images were cropped from Movie 6.

### Mice lacking *Kif6* or *Kif9* develop hydrocephalus and exhibit male infertility

To uncover their physiological functions, we generated *Kif6* and *Kif9* knockout mice (Fig. 5a, b) using the CRISPR/Cas9 system (Joung et al., 2017). Immunoblotting confirmed complete depletion of Kif6 or Kif9 in motile cilia-enriched tissues (Fig. 5c). Both *Kif6^-/-^* and *Kif9^-/-^* mice were born at the expected Mendelian ratio of genotypes. The majority of *Kif6^-/-^* mice experienced growth failure and often exhibited a dome-shaped skull compared to wild-type littermates after P21, suggestive of severe hydrocephalus (Fig. 5d, arrows). Some of these mice also displayed a forward curvature of the spine and a smaller size compared to their littermates (Fig. 5d, arrowhead). Careful examination of brain dissections or coronal sections revealed that every analyzed *Kif6^-/-^*mouse (100%; total n = 60) displayed lateral ventricle expansion when compared to the wild-type controls (Fig. 5f, h). This ventricle enlargement was evident even in mice that displayed a normal skull appearance and longer lifespan (Fig. 5f, bottom). In sharp contrast to *Kif6^-/-^* mice, *Kif9^-/-^*mice appeared normal similar to their wild-type littermates based on macroscopic examination (Fig. 5e). However, coronal brain sections revealed enlarged ventricles in over half of *Kif9^-/-^* mice (Fig. 5g, h), indicative of hydrocephalus. Survival curve analysis revealed that approximately 77% of *Kif6^-/-^* mice died before reaching P90, while *Kif9^-/-^* mice exhibited a normal lifespan similar to their wild-type littermates during a 12-month observation period (Fig. 5i).

**Figure 5.**
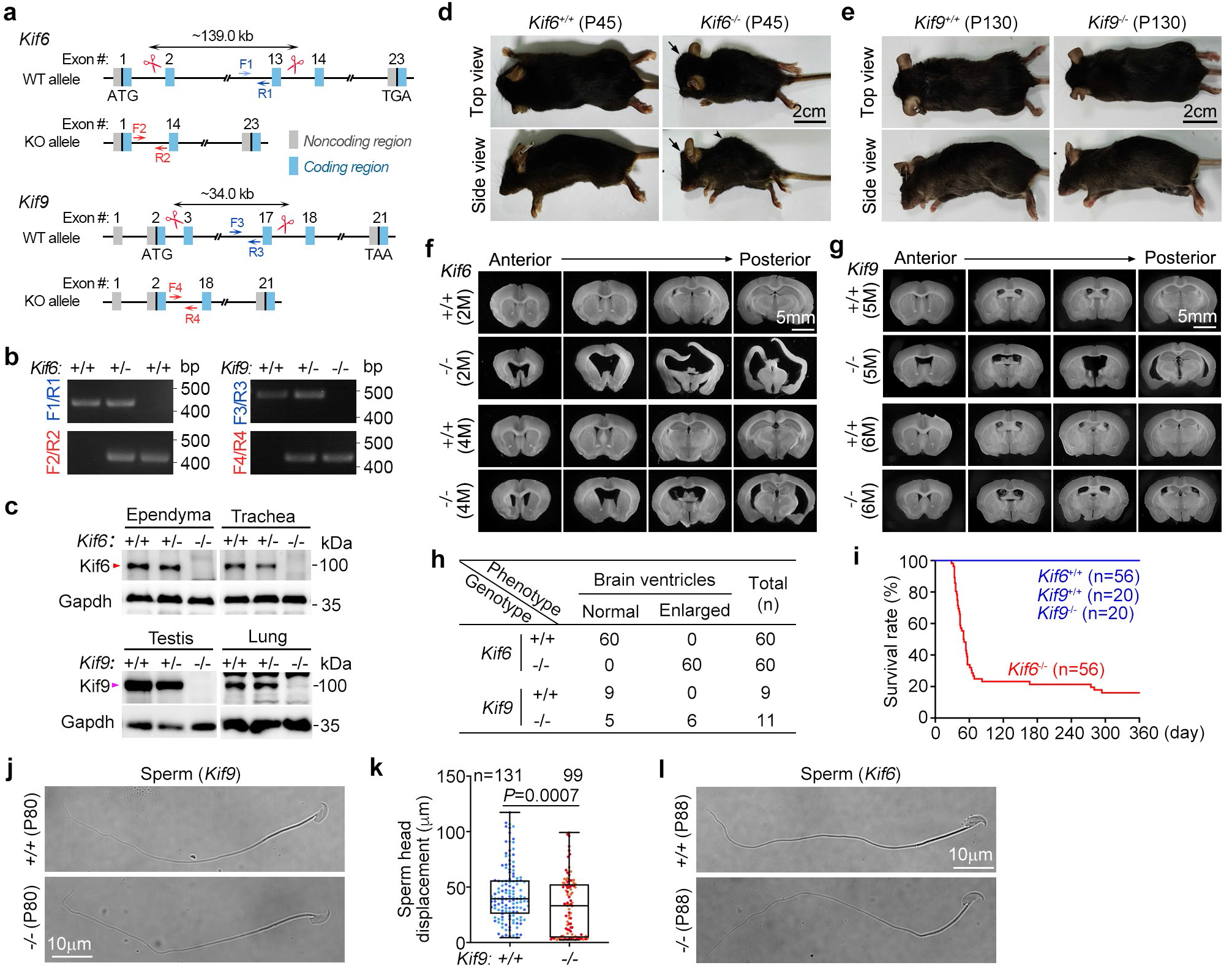
Knockout of *Kif6* or *Kif9* leads to hydrocephalus and male infertility. **(a)** Schematic for generating *Kif6* (top) and *Kif9* (bottom) knockout mice using CRISPR/Cas9 technique. The targeting of sgRNA (scissors) and genotyping PCR primers were indicated for each knockout model. **(b)** Representative genotyping PCR results. Genomic DNAs were extracted from P21 mice tails and amplified by PCR using the indicated primers in (**a**). **(c)** Complete depletion of Kif6 and Kif9 proteins in the indicated tissues. Tissue lysates were prepared from 2-month-old mice of the indicated genotypes. Gapdh served as a loading control. **(d, e)** Morphologies of *Kif6* and *Kif9* deficient mice. *Kif6*^-/-^ mice exhibited a dome-shaped skull (arrow) and kyphosis (arrowhead) with skinny bodies (**d**). In contrast, *Kif9^-/-^* mice showed a normal appearance (**e**). **(f, g)** Representative images of coronal brain sections of *Kif6*^-/-^ (**f**) or *Kif9^-/-^* (**g**) and their corresponding wide-type littermates. M: month. **(h)** Quantification of mice with enlarged brain ventricles (n = 60 for *Kif6*^-/-^; n = 11 for *Kif9^-/-^*mice). **(i)** Survival curves of mice in indicated genotypes during a 12-month observation period. The number (n) of mice used for each genotype were indicated. **(j)** Sperm morphology from wide-type and *Kif9*^-/-^ littermates. **(k)** *Kif9* deficiency exhibited sperm forward motility defects. Quantification results of sperm head displacement (μm) within 1 second from two mice each genotype (10 movies per mouse) were pooled together (mean ± SD plus sample dots) and subjected to unpaired two-tailed student’s *t*-test. Two pairs of P80 *Kif9*^-/-^ mice and wild-type liitermates were used. The indicated sperm numbers (n) were presented. **(l)** Sperm morphology from wide-type and *Kif6*^-/-^ littermates. Two pairs of P88 *Kif6*^-/-^ mice and wild-type litermates were used.

While female *Kif9^-/-^* and *Kif6^-/-^* mice were fertile, *Kif9^-/-^* and *Kif6^-/-^* males were sterile, even when those with normal appearance were used for mating (n = 3 for each genotype). Subsequently, we isolated sperms from the cauda epididymis of wild-type, *Kif9^-/-^* and *Kif6^-/-^* mice at reproductive age (∼ 2 months) for motility analysis. *Kif9^-/-^* male mice showed a slight decrease in sperm count compared to wild-type controls (Supplementary Fig. 4a, b). Although *Kif9^-/-^* sperms had normal morphologies, they displayed impaired progressive motilities (Fig. 5j, k and Movie 7), consistent with previous report (Miyata et al., 2020). *Kif6^-/-^* mice, however, showed a significant reduction in sperm count (Supplementary Fig. 4c, d). Interestingly, while *Kif6^-/-^* sperms appeared to be normal in morphologies (Fig. 5k), the majority of them completely lacked motilities (67%, total n = 46) as compared to wild-type sperms (3%, n = 131) (Movie 8). In contrast to these, we did not observed noticeable defects, such as polydactyly or polycystic kidney, in organs where primary cilia are critical (Nachury and Mick, 2019), suggesting intact primary cilia functions. These observations emphasize the critical yet distinct roles of Kif6 and Kif9 in motile cilia and sperm flagella.

### *Kif6* but not *Kif9* deficiency disrupts the planar polarity of ependymal motile cilia

The planar beating of ependymal multicilia in mice older than P21 generates unidirectional CSF flow in brain ventricles (Guirao et al., 2010; Ohata and Alvarez-Buylla, 2016). As abnormalities in mouse ependymal cilia cause hydrocephalus (Ji et al., 2022; Liu et al., 2021; Ohata et al., 2014), we dissected the brain of >P21 mice and stained multicilia in living ependymal tissues with SiR-tubulin, a fluorescent probe for MTs (Lukinavicius et al., 2014), followed by live imaging using high-speed spinning disk confocal microscopy (Fig. 6a). We did not observe gross differences in multiciliogenesis, the back-and forth beat pattern of multicilia, and their beat frequency between *Kif6^+/+^* and *Kif6^-/-^*or *Kif9^+/+^* and *Kif9^-/-^* littermates (Fig. 6b-e and Movies 9, 10). Nevertheless, while multicilia beat directionally across different cells in the wild-type or *Kif9^-/-^* ependymal tissues, their beat directions differed among individual cells in *Kif6^-/-^*ependymal tissues regardless of the hydrocephalus severity of the mice (Fig. 6f, g and Movies 9, 10), indicating a disruption in the planar polarity of *Kif6^-/-^*multicilia (Ohata and Alvarez-Buylla, 2016).

**Figure 6.**
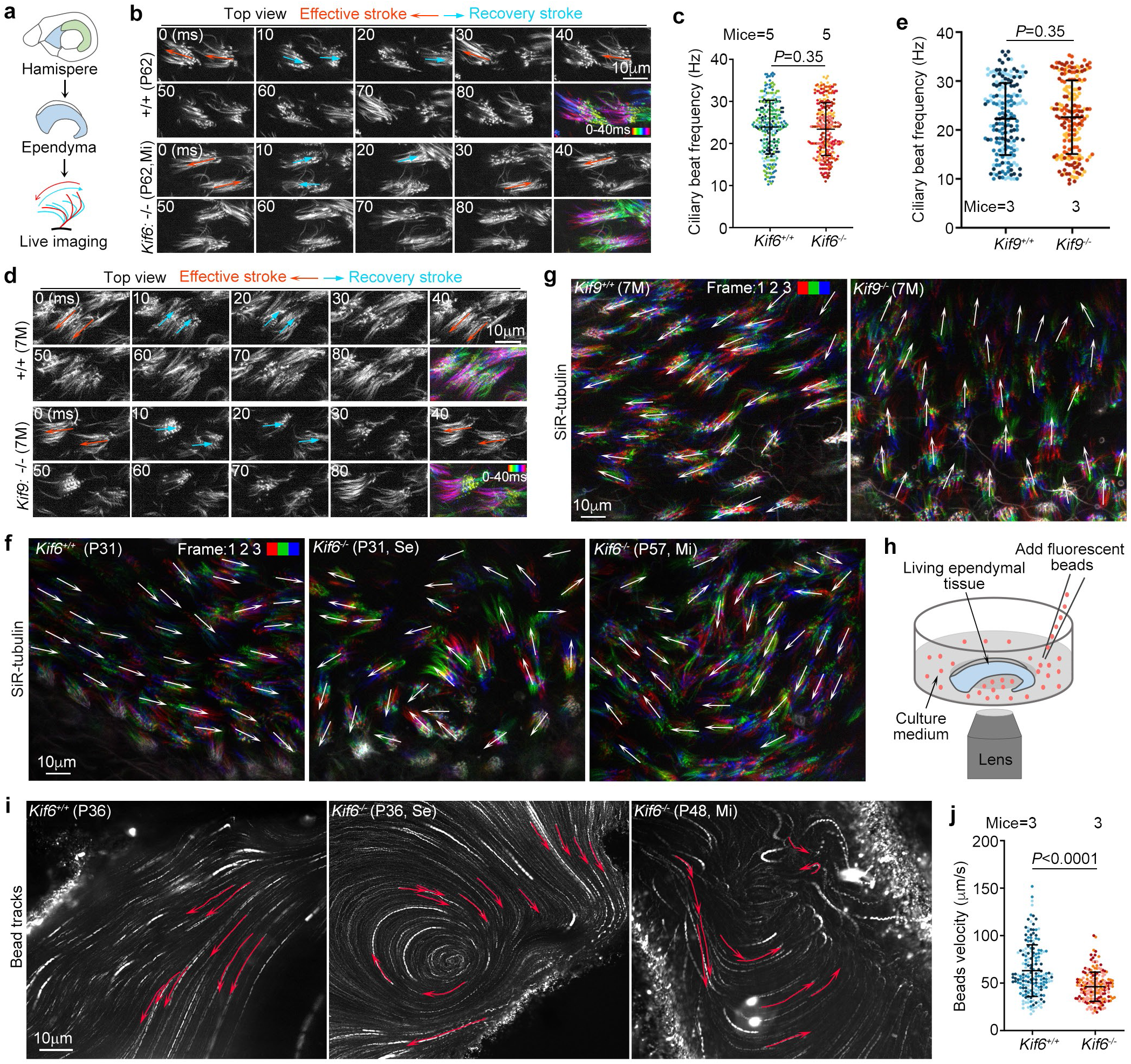
Knockout of *Kif6* impairs planar polarity of cilia across the ependyma. Quantification results are pooled together (mean ± SD plus sample dots) and subjected to unpaired two-tailed student’s *t*-test. Mi: mild hydrocephalus mice with normal appearance but larged ventricles; Se: severe hydrocephalus mice with the observable dome-shaped skull like Figure 1d. **(a)** Schematic of experimental design. Whole-mounts of ventricle walls were freshly dissected and subjected to electron microscopy (EM) or live cell imaging. SiR-tubulin, a fluorescent dye for MTs, was used to label cilia. Ciliary motility was live-cell imaged at a single optical section with 10-ms intervals. An illustration of the beat pattern of an ependymal cilium, depicted as a side view, is provided for better comprehension. **(b-e)** The directional ciliary beat in ependymal cells and ciliary beat frequencies of the indicated genotypes. No gross differences were observed in the back-and forth beat pattern of multicilia (**b**, **d**) and their beat frequencies (**c**, **e**) between *Kif6^+/+^*and *Kif6^-/-^* or *Kif9^+/+^* and *Kif9^-/-^*littermates. Two consecutive cycles of cilia beat patterns were showed and color coded arrows were used to indicate the patterns. The first cycle of cilia beating was pseudo-colored and merged (**b**, **d**). Quantification of ciliary beating frequencies (**c**, **e**). Results were from 5 pairs of littermates with 60 cells per mouse for (**c**) or 3 pairs of littermates with 50 cells per mouse for (**e**). **(f)** Unidirectional multicilia beat across ependyma was disrupted by the loss of *Kif6*. The first three consecutive frames cropped from Movies 9-10, were pseudo-colored and overlaid to show ciliary motilities. Beating directions of multicilia are indicated by arrows. **(g)** Planar polarity of cilia beating was not altered in *Kif9*-deficient ependyma. **(h)** Schematic illustrations forthe beads assay. Fluorescent beads, suspended in culture medium, were imaged in areas covered with ependymal tissues. The imaging was performed at a fixed z-plane, with an exposure time of 100 ms per frame to record the motilities of the beads. **(i)** Representative projections of bead movements from Movies 11-12. Trajectories of traceable, rapidly-moving beads in the first 500 ms were superimposed to show flow directions. **(j)** Quantification of bead velocities. Results from two pairs of P36 and one pair of P48 littermates, with 60 traceable beads for each.

Next, we investigated whether the loss of planar beating impaired directional fluid flow across *Kif6^-/-^* ependymal tissues by adding latex fluorescent microbeads to the culture medium and tracking their movements by live imaging (Fig. 6h). We observed that the beads underneath wide-type ependymal tissues flew in one direction with a velocity of 63.6 ± 27.1 μm/s (Fig. 6i, j and Movie 11). In contrast, the beads underneath *Kif6^-/-^* ependymal tissues mainly oscillated or whirled regionally, with a velocity of 46.3 ± 15.4 μm/s (Fig. 6i, j and Movie 12). Therefore, we attribute the hydrocephalus of *Kif6^-/-^* mice (Fig. 6) to impaired CSF flow.

### Kif6 is crucial for the rotational polarity of BBs in ependymal tissues

For insights into why *Kif6^-/-^* multicilia lost the planar polarity of beat directions (Fig. 6), we performed transmission electron microscopy with ependymal tissues to examine cilia-related ultrastructure (Fig. 7a). Cross-sections indicated that *Kif6^-/-^* ependymal cilia still contained 9+2 axonemes, with outer and inner dynein arms on DMTs (Fig. 7b). Nevertheless, the image resolutions were still insufficient for a definite conclusion whether the entire axonemal ultrastructure was the same as the wild-type one (Fig. 7b) (Meng et al., 2023). Interestingly, we observed that, while BFs were oriented towards a similar direction in the wild-type ependymal cells, a state termed the rotational polarity of BBs (Ohata and Alvarez-Buylla, 2016), BFs in *Kif6*^-/-^ ependymal cells were misoriented regardless of the severity of hydrocephalus (Fig. 7c). To quantify the extent of the rotational polarity, we considered each arrow marking the BF orientation as a unit vector and measured the mean vector length for each electron micrograph (Bustamante-Marin et al., 2019). The proper rotational polarity of BBs in the wild-type ependymal tissues was confirmed by a mean vector length of 0.97 ± 0.07. In sharp contrast, the value was 0.35 ± 0.17 and 0.31 ± 0.18 for ependymal tissues from mice respectively with mild and severe hydrocephalus (Fig. 7d).

**Figure 7.**
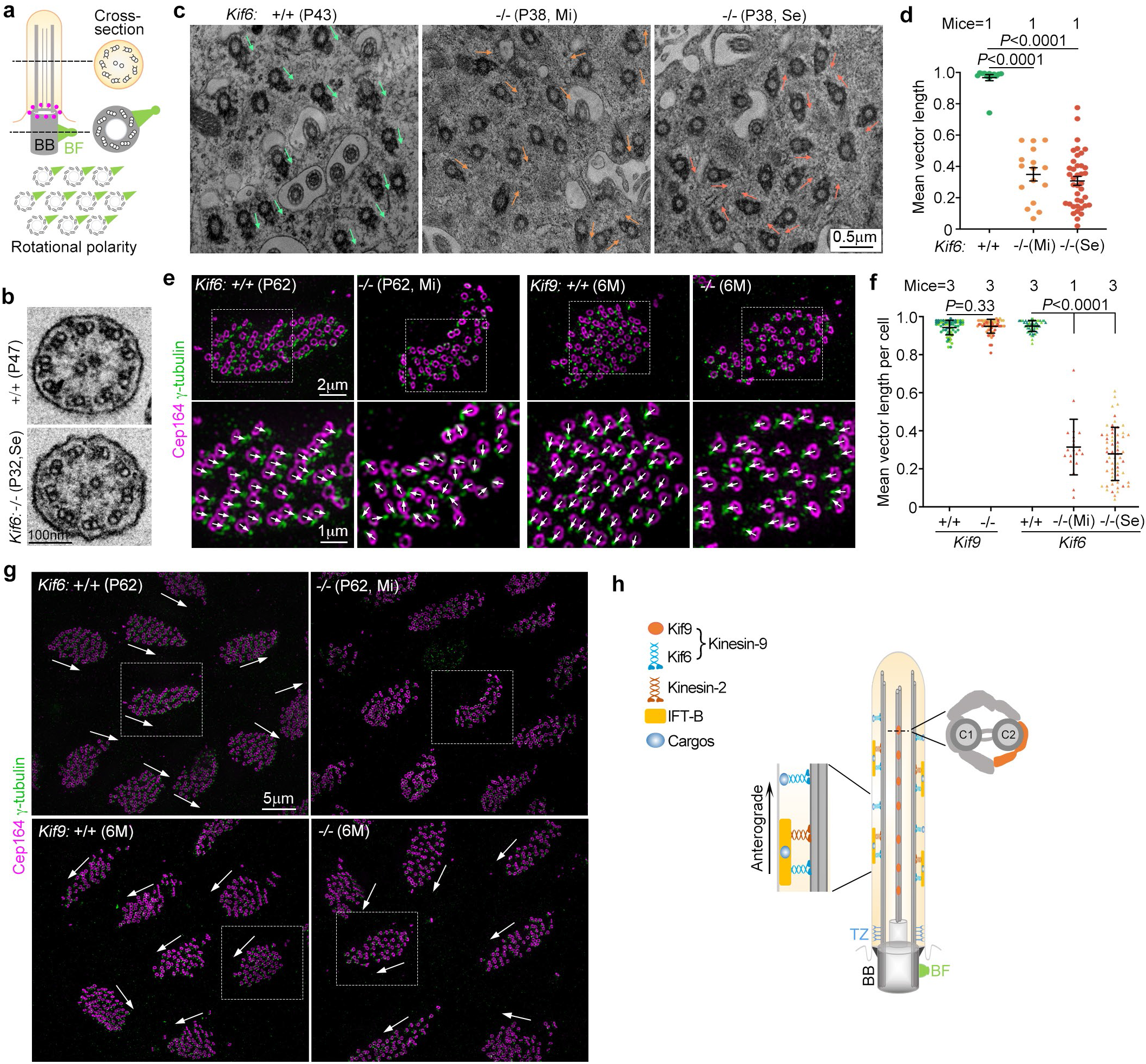
*Kif6* deficiency impairs rotational polarity of BBs in ependymal tissues. Quantification results are pooled together (mean ± SD plus sample dots) and subjected to unpaired two-tailed student’s *t*-test. Se: Severe hydrocephalus mice with obvious dome-shaped skull. Mi: mild hydrocephalus mice with normal morphological appearance but enlarged ventricles. **(a)** Schematic diagram of a motile cilium and an uniformed rotational polarity of BBs. A side view of a motile cilium and transverse views of the axoneme and the BB were presented. BB polarity is denoted by their conical BFs that extend laterally to the MT triplets. **(b)** Representative transmission electron microscopy images showed transverse views of the axoneme of ependymal cilia from wild-type and *Kif6^-/-^* mice (n = 2 biological replicates). **(c, d)** Representative transmission electron microscopy images (**c**) and quantification analysis on the extent of rotational polarity (**d**). BF orientations are indicated by arrows. The mean vector length (**d**) was calculated from BBs with clear BFs in at least ten full-size electron micrographs and over 130 BFs were scored for each genotype. **(e-g)** The loss of *Kif6*, but not *Kif9*, impaired rotational polarity. Ependymal tissues from the indicated mice were immunostained for Cep164 to visualize BBs and γ-tubulin to visualize BFs. Framed regions (**e**) were magnified to show the rotational polarity of BBs in individual cells **(g)**. The BF orientations are indicated by arrows. Note the misorangized rotational polarity in both cellular and tissue-level in *Kif6^-/-^*ependyma. Mean vector lengths (**f**) were scored from 20 cells each mouse. Mice used were two 6-month-old (6M) and one 7-month-old (7M) mice for *Kif9^-/-^* and three of their wild-type littermates; two P34, one P43 mice with severe hydrocephalus and one P62 mouse with mild hydrocephalus for *Kif6^-/-^* and their corresponding wild-type littermates. **(h)** A summary model. Kif6 acts as a motile cilia-specific IFT motor, either on its own or in association with the IFT-B-containing IFT train. On the other hand, Kif9 localizes on the central apparatus to modulate detailed ciliary beat forms, possibly by inducing conformational changes in the central apparatus as a C2 projection (Han et al., 2022). BB, basal body; BF, basal foot; TZ, transition zone.

As transmission electron microscopy is not ideal for assessing BB polarities in entire cells, letting alone planar polarities, we immunostained whole-mounts of ependyma from wild type, *Kif6^-/-^*, or *Kif9^-/-^* mice using Cep164, a transition fiber component (Siller et al., 2017), to mark BBs, and γ-tubulin or Centriolin to label BFs (Nguyen et al., 2020; Ohata et al., 2014). 3D-SIM revealed a uniform rotational polarity of BBs in individual cells and a planar polarity among neighboring cells in both wild type and *Kif9^-/-^*wholemounts (Fig. 7e-g and Supplementary Fig. 5a-c). Both the rotational and the planar polarities of BBs, however, were markedly impaired in *Kif6^-/-^* ependymas (Fig. 7e-g and Supplementary Fig. 5a-c). We also scored BB numbers and observed that the deficiency of either *Kif6* or *Kif9* did not interfere with the BB biogenesis (Supplementary Fig. 5d), consistent with the normal ciliogenesis (Fig. 6) but different from the reported defective ciliogenesis in *Kif6^p.G555fs^* mice (Konjikusic et al., 2018).

In addition to the rotational polarity, BBs in ependymal cells form patches that tend to be positioned towards one side of the cell border, a state termed the translational polarity (Mirzadeh et al., 2010; Ohata and Alvarez-Buylla, 2016). We quantified the extent of BB patch displacement relative to the cell center (Mirzadeh et al., 2010) but did not observe a significant change between the wild-type mice and their *Kif6^-/-^*littermates (Supplementary Fig. 5e-g). The area of BB patches was also unaffected in *Kif6^-/-^* mice with mild hydrocephalus but slightly increased in mice with severe hydrocephalus (Supplementary Fig. 5e, f, h), possibly due to the pathological effects of the hydrocephalus.

Taken together, we conclude that Kif6 is critical for the rotational polarity of BBs but dispensable for the transitional polarity. The defective rotational polarization in *Kif6^-/-^* ependymas abolishes the planar polarity of ciliary beat, leading to abnormal CSF flow and hydrocephalus.

## Discussion

In this study, we elucidated the distinct ciliary localizations and functions of Kif6 and Kif9, two kinesin-9 family members in mammalian motile cilia (Fig. 1b and 7h). Kif9 is mainly found in the CP lumen in motile cilia, while Kif6 as puncta localizes and moves bidirectionally with or without IFT-B particles along the DMTs (Figs. 1-3 and Supplementary Figs. 1-2). Interestingly, neither Kif6 nor Kif9 are necessary for ciliogenesis (Fig. 6). However, *Kif6* knockout disrupts cilia motility coordination (Fig. 6 and Movie 10) and impaired sperm motility (Fig. 5l and Movie 8), and leads to hydrocephalus (Fig. 4d, f, h) and male infertility in mice. *Kif9* knockout results in male infertility by affecting sperm forward motility (Fig. 5k and Movie 7) and a mild hydrocephalus (Fig. 5g, h) possibly due to a defect in fine-tuning the cilia beating.

IFT is responsible for bidirectional transportation of ciliary proteins. The kinesin-2 family motors, including Kif3a/3b/3c and Kif17, facilitate the anterograde IFT, which is essential for ciliogenesis of both primary and motile cilia across species (Klena and Pigino, 2022; Scholey, 2013; Zhao et al., 2012). Compared to IFT in primary cilia, IFT in motile cilia transports a wide variety of large protein complexes, including dynein arms, radial spokes, and proteins critical for cilia beating. Futhermore, the axonemal MTs of motile cilia are frequently decorated with an array of associated proteins or protein complexes and exhibit rapid beating (Gui et al., 2021; Han et al., 2022; Ma et al., 2019). These differences suggest that IFT in motile cilia may possess unique features or require additional molecular motors. Remarkably, using high-speed and super-resolution GI-SIM, we performed the first dual-colour imaging of particles trafficking within mammalian motile cilia. We discovered that Kif6, but not Kif9, undergoes processive movement in ependymal cilia (Figs. 2-3 and Supplementary Fig. 2; Movies 2-5). The bidirectional movement of Kif6 along the ciliary axoneme, either with or without IFT-B particles (Fig. 3 and Supplementary Fig. 2d), together with its ability to bind and glide along MTs, implies that Kif6 may transport cargos either independently or in association with the IFT-B-containing IFT train (Fig. 7h). This finding suggests the possibility of separate protein transport mechanisms, warranting further investigation. It is important to note that the presence of endogenous proteins can not currently be ruled out in IFT-B free Kif6 puncta. The presence of Kif6 in single-cell organisms such as *Chlamydomonas* and *Trypanosoma*, which contain motile cilia/flagella (Bernstein et al., 1994; Demonchy et al., 2009), suggest its conserved role in IFT.

In *Trypanosoma*, the lack of *KIF9B*, a *Kif6* homolog, disrupts the assembly of the PFR, a structure asymmetrically assembled at DMTs 4-7 along the flagellar axoneme, required for flagellar motility (Demonchy et al., 2009; Zhang et al., 2021). While the PFR is exclusive to euglenoids and kinetoplastids (Cachon et al., 1988; Langousis and Hill, 2014), the “9+2” mammalian motile cilia also display asymmetrical structures enabling the back-and-forth ciliary beat (Ishikawa, 2017; Loreng and Smith, 2017). The ciliary beat direction is perpendicular to the CP plane from DMT 1 to the gap between DMT 5 and 6 (axonemal orientations), and the direction of BFs aligns with the effective stroke of cilia (BB polarities) (Butler and Wallingford, 2017; Clare et al., 2014; Lindemann and Lesich, 2021; Ohata and Alvarez-Buylla, 2016). Our recent study reveal that axonemal orientations and BB polarities are initially decoupled and gradually become aligned during the development of multiciliated ependyma (Pan et al., 2023). This process requires both the PCP pathway and external flow force. *Kif6* deficiency does not affect translational polarity of BBs, but it does disrupt the rotational polarity of BBs and planar polarity of cilia in ependyma (Figs. 6-7 and Supplementary Fig. 5; Movie 10). Since PCP pathway is key for translational polarity (Mirzadeh et al., 2010; Ohata and Alvarez-Buylla, 2016), Kif6 likely does not function upstream of the PCP pathway. Instead, it may transport specific cargos containing factors important for hydrodynamic force responses or the axoneme-BF coupling (Fig. 7h) (Guirao et al., 2010; Pan et al., 2023; Tissir et al., 2010). This idea is supported by the misalignment of BBs in *Kif6^-/-^* mice, similar to obervations in cultured mEPCs lacking external flow force (Guirao et al., 2010). Furthermore, this speculation is also reinforced by the observation that *Kif6^-/-^* mice do not exhibit noticeable airway defects. The rotational polarity of BBs in tracheal epithelial cells can align correctly in the absence of flow forces *in vitro* (Herawati et al., 2016). In the future, identifying molecules or factors associated with Kif6 could provide more insights into its function and the regulatory mechanism.

The immobilization of Kif9 within ependymal cilia is consistent with the observation in *Xenopus* epidermal multiciliated cells (Konjikusic et al., 2023), reinforcing the notion that Kif9 is not involved in ciliary transport. In flagellated species *Trypanosoma* and *Chlamydomonas*, knockdown *Kif9* homolog, lead to a reduction flagella movement (Bernstein et al., 1994; Demonchy et al., 2009; Yokoyama et al., 2004). High-resolution structures of *Chlamydomonas* by cryo-electron microscopy (Cryo-EM) shows that the Kif9 homolog, KLP1, exhibits an 8-nm sliding on the C2 MT and is proposed to function as an active motor to power the movement of the central apparatus for regulating flagella beating (Han et al., 2022). Considering the conserved CP localization from protists to mammals and defective sperm motility and hydrocephalus upon its depletion in mice (Fig. 5g, h, k and Movie 7), it could be hypothesized that mammalian Kif9 as a central apparatus regulator for cilia/flagella beating, through mediating conformation changes of central apparatus during beating (Fig. 7h). Future studies employing advanced technologies such as cryo-electron tomography technology (Cryo-ET) studies will be needed to validate this hypothesis.

## Methods

### Plasmids

The cDNAs encoding full-length *Kif6* (GenBank: NM_177052.3), *Kif9* (GenBank: NM_001163569.1), *Ift81* (GenBank: NM_001358917.1), and *Ift27* (GenBank: NM_025931.3) were amplified using PCR with cDNAs extracted from cultured mEPCs (Day 10) as the template. To generate adenoviral expression constructs, the cDNAs fragments were PCR-amplified and subsequently cloned into the entry vector pYr-1.1-GFP (YRBio, China) or pYr-1.1-Halo to express GFP-tagged or Halo-tagged proteins, respectively. The LR recombination reactions between the entry constructs and the destination vector pAd/BLOCK-iT-DEST (YRBio, China) were carried out using LR Clonase II enzyme mix (Thermo Fisher, 11791020), as described previously (Zhao et al., 2021).

The rat Kif5c-MD-GFP (1-430 aa) construct was kindly provided by Prof. Bao (Institute of Biochemistry and Cell Biology, CAS). To express GFP-fusion proteins, the cDNAs of Kif6 (1-802 aa), Kif6-MD (1-353 aa), Kif6-CC1 (1-385 aa), Kif6-CC2 (1-494 aa), Kif9 (1-810 aa), Kif9-MD (1-348aa), Kif9-CC1 (1-442 aa) and Kif9-CC2 (1-694 aa) were amplified and subcloned into pEGFP-N1. To express GFP-Flag-fusion proteins, the cDNAs of Kif6-GFP, Kif6-MD-GFP, Kif6-CC1-GFP, Kif6-CC2-GFP, Kif9-GFP, Kif9-MD-GFP, Kif9-CC1-GFP, Kif9-CC2-GFP and Kif5c-MD-GFP were amplified and constructed into pCDAN3.1-CFlag. To generate polyclonal antibodies of Kif9, the cDNA fragments encoding the indicated amino acids (348-810 aa) were amplified by PCR and subcloned into pET-28a or pGEX-4T-1, to express His-tagged or GST-tagged Kif9, respectively.

All constructs were validated by sequencing. The primers used for PCR amplification are listed in Supplementary Table 1.

### Mice

All animal experiments were performed following guidelines approved by the Institutional Animal Care and Use Committee of Institute of Biochemistry and Cell Biology. The mice used in this study had a genetic background of C57BL/6J. *Kif6^+/-^* and *Kif9^+/-^* mice were obtained from GemPharmatech, China. Ages of mice used in this study were indicated in the figures or the legends. gRNA sequences used for targeting *Kif6* or *Kif9* and primers used for genotyping are listed in Supplementary Table 1.

### Cell culture and transfection

HEK293T and HEK293A cells were grown in Dulbecco’s modified Eagle’s medium (DMEM) (Thermo Fisher, 12430-054) supplemented with 10% fetal bovine serum (FBS) (Ausbian, VS500T), 0.3 mg/ml glutamine (Sigma, G8540), 100 U/ml penicillin (Solarbio, P8420) and 100 U/ml streptomycin (Solarbio, S8290). NIH3T3 cells were cultured in DMEM medium with 10% normal goat serum (NGS), 0.3 mg/ml glutamine, 100 U/ml penicillin, and 100 U/ml streptomycin. hTERT-RPE1 and IMCD3 cells were cultured in DMEM/F12 medium (GE Healthcare) supplemented with 10% FBS, 0.3 mg/ml glutamine, 100 U/ml penicillin, and 100 U/ml streptomycin. Additionally, RPE1 culture medium was supplemented with 10 μg/ml hygromycin B (Thermo Fisher). All cell lines were passaged at a 1:4 ratio once they reached 80%-90% confluency. HEK293A cells were used for packaging adenovirus expressing GFP-tagged proteins following the guidelines (YRBio, China). All the cells were routinely tested for mycoplasma contamination.

Primary culture of mouse ependymal cells (mEPCs) was prepared and cultured as previously described (Delgehyr et al., 2015; Zhao et al., 2021). Newborn C57BL/6J mice were anesthetized on ice, and the cerebellum, olfactory bulbs, meninges, and hippocampus were carefully removed using sharp tweezers (Dumont, 1214Y84) in cold dissection solution (5 mM Hepes, 161 mM NaCl, 5 mM KCl, 1 mM MgSO_4_, 3.7 mM CaCl_2_, and 5.5 mM Glucose, pH 7.4) under a stereo microscope. The remaining telencephala were then digested in a dissection solution containing 10 U/ml papain (Worthington-Biochem, LS003126), 0.2 mg/ml L-cysteine, 1.5 mM NaOH, 1 mM CaCl_2_, 0.5 mM EDTA, and 0.15% DNase I (Sigma-Aldrich, D5025) for 30 min at 37°C. After gentle pipetting and centrifugation at 1,500 rpm for 5 min at room temperature (r.t.), the cells were resuspended in DMEM medium supplemented with 10% FBS and 1% penicillin/streptomycin and seeded into flasks (Sigma-Aldrich, L2020) pre-coated with 10 μg/ml Human Plasma Fibronectin (Sigma-Aldrich, FC010). Neurons were mechanically shaken off and removed by patting the flask approximately 2 days after inoculation. When reaching ∼90% confluence, cells were digested with trypsin and re-plated into fibronectin-coated 29-mm glass-bottomed dishes (Cellvis, D29-14-1.5-N) for immunofluorescence staining. Once cells were cultured to ∼90% confluency, FBS was removed from the medium to induce differentiation.

For the primary culture of mTECs, we followed the method described (Zhao et al., 2021). mTECs were used to investigate the expression profile of *Kif6* and *Kif9* during the differentiation of mTECs, because of their high induced differentiation efficiency.

For transfection of plasmids, cells were transfected at ∼70% confluency with Lipofectamine 2,000 (Thermo Fisher).

### Preparation and culture of whole-mounts of lateral ventricle walls

Whole-mounts of the lateral ventricles were prepared as described (Ohata et al., 2014) with minor modifications. Mice were euthanized with CO_2_, and the lateral ventricles were carefully dissected at a thickness of 500-1000 μm using Vannas Scissors (66VT, 54140B) in pre-warmed (37°C) dissection solution (25 mM Hepes,117.2 mM NaCl, 5.3 mM KCl, 0.81 mM MgSO_4_, 1.8 mM CaCl_2_, 26.1 mM NaHCO_3_, 1 mM NaH_2_PO_4_·2H_2_O, and 5.6 mM Glucose, pH 7.4). The samples were subsequently cultured in DMEM supplemented with 0.3 mg/ml glutamine, 100 U/ml penicillin, and 100 U/ml streptomycin.

### Viral production and transfection

Adenovirus particles were utilized to infect mEPCs or IMCD3 cells and produced in HEK293A cells as previously described (Zhao et al., 2021). In brief, the adenovirus expression constructs were linearized using *P*acI restriction enzyme (NEB) and transfected into HEK293A cells cultured in 6-well plates using Lipofectamine 2,000 to produce the initial adenovirus. When approximately 80% of the cells exhibited a cytopathic effect (CPE), the culture medium was collected and infected HEK293A cells in a 10-cm dish. Once about 80% of the cells showed a CPE, the culture medium was collected to infect fresh HEK293A cells for viral amplification. When ∼80% of the cells exhibited a CPE, both cells and the culture medium were collected. To ensure complete release of the viral particles, three freeze-thaw cycles (-80°C and 37°C) were performed, followed by centrifugation at 1,500 rpm for 5 min to remove cell debris. The harvested adenoviral particles were used at a dilution of 1:1000 or 1:200.

### Immunofluorescence staining and imaging

IMCD3 cells were fixed with 4% paraformaldehyde (PFA) in PBS, whereas mEPCs grown on 29 mm glass-bottom dishes and freshly dissected whole-mounts were pre-permeabilized with 0.5% Triton X-100 in PBS for 30 sec before fixation to remove soluble proteins. After fixation, the cells were extracted with 0.5% Triton X-100 in PBS for 15 min and blocked in 4% BSA in TBST at r.t. for 1 h. Next, the cells were incubated with primary antibodies overnight at 4°C. After three washes with the blocking solution for 5 min each, the cells were incubated with secondary antibodies at r.t. for 1h, and mounted in Dako (Dako, S3023) or ProLong^TM^ (Thermo Fisher, 2273640). All the antibodies used are listed in Supplementary Table 2.

Fluorescent images were captured using a Leica TCS SP8 WLL system with a HC PL APO CS2 63 × /1.40 OIL objective or an Olympus Xplore SpinSR10 microscope with UPLAPO OHR 60 ×/1.50 Objective Lens, and images were processed with maximum intensity projections. 3D-SIM super-resolution images were taken with a DeltaVision OMX SR system (GE Healthcare) and the z-axis scanning step was 0.125 μm. The original images were processed for OMX SI Reconstruction, OMX Image Registration, and maximum intensity projection with SoftWoRx software.

To perform GI-SIM imaging for mEPCs, the cells were seeded onto fibronectin-coated 12-mm coverslips and were infected with adenovirus one day prior to serum starvation. To fluorescently label their multicilia, mEPCs were incubated with 100 nM SiR-tubulin (Spirochrome, SC002) for 30 min or 200 nM Janelia Fluor® 549 HaloTag ligand (Promega, GA1110) for 20 min. After labeling, mEPCs were gently laid flat against the bottom coverslip and GI-SIM imaging was performed as described (Guo et al., 2018; Qiao et al., 2022). For dual-color imaging, instrument calibration was conducted routinely with fluorescent beads. The images and movies were processed using Fiji and Huygens softwares. The velocities and frequencies of GFP or Halo-tagged particles were manually analyzed by kymographs with Fiji software as previously described (Lechtreck, 2013). For analyzing particle frequencies, we only used the first 200 frames of a recording to avoid potential bleaching of the signal over the time course of image capture.

To record ciliary beating, the whole mounts of lateral ventricles were dissected and cut at a thickness of 500-1000 μm as previously described (Ohata et al., 2014; Zhao et al., 2021). To label the cilia, 100 nM SiR-tubulin was added to the culture medium and incubated for 1 h. High-speed live imaging was conducted at 37℃ using an Olympus Xplore SpinSR 10 microscope equipped with UPLAPO OHR 60 × /1.50 Objective Lens, Hamamatsu ORCA-Fusion camera, 4,000 rpm CSU Disk Speed, and OBIS Laser. Ciliary beat was captured at 100 frames per second (fps) by a time-lapse collection of single optical sections with 80% laser power (640 nm) to achieve an exposure time of 5 ms. Kymographs were generated for the first 500 ms using Fiji.

The ependymal flow assay was performed as described (Liu et al., 2021; Mirzadeh et al., 2010) with minor modifications. Briefly, the freshly dissected ependyma was sliced using a vibratome (Leica VT1000S) to obtain slices with a thickness of 250 μm. Before imaging, multicilia were labeled with 100 nM SiR-tubulin for 1 h and small fluorescent latex beads with a diameter of 250 nm at 1:500 dilution (Invitrogen, F8809) were added to the culture medium. Time-lapse microscopy was performed at 37 ℃ using using an Olympus Xplore SpinSR10 microscope equipped with UPLAPO OHR 60 × /1.50 Objective Lens. The laser power for the beads channel (561 nm) was adjusted based on the fluorescent intensity, and exposure time was set to 100 ms. The laser power for the SiR-tubulin channel (640 nm) was set to 80% to achieve an exposure time of 5 ms.

During recording, fluorescent beads suspended in the culture medium containing the living ependymal tissues were imaged at a fixed z-plane to record their motility. Subsequently, multicilia in the same areas were visualized by adjusting the focal plane, and live imaging of a single optical section was conducted to record ciliary motility. All frames of beads in the same area were projected together from Movies 12-13 and superimposed with trajectories of traceable, rapidly-moving bead aggregates in the first 500 ms to show flow directions and tracks. The velocity of beads was meausured using Fiji.

To record sperm motility, sperm were released from cauda epididymidis into buffer (Njabsw, M1130), and followed by live imaging with a spinning disk confocal microscope at a fixed z-plane and at 25-ms intervals. Sperms from two pairs of P88 wild-type and *Kif6*^-/-^ litermates and two pairs of P80 of wild-type and *Kif9*^-/-^ litermates littermates were examined. For analyzing sperm number, we randomly select the first frame (full micrograph with 2304 × 2304 pixels) of a recording and only count the sperm with head from each genotype. For analyzing sperm head displacement, we randomly select a recoding last for 1 second to trace the clear sperm movement and then measure the head displacement.

### Protein purification

Protein purification was performed as described (Liu et al., 2021). Cells were lysed with cold lysis buffer (20 mM Tris-HCl, pH 7.5, 100 mM KCl, 0.1% NP-40, 1 mM EDTA, 10 mM Na_4_O_7_P_2_, 10% Glycerol, and protease inhibitors (Sigma, 539134)). The lysates were collected after centrifugation for 20 min at 13,000 rpm at 4℃ to remove debris. The precleared lysates were incubated with 20 μl of anti-FLAG beads (Sigma, A2220) for 2 hr at 4℃ under rotary agitation. The beads were washed three times with the lysis buffer and three times with wash buffer (20 mM TrisHCl, pH 7.5, 150 mM KCl, 0.5% NP-40, 1 mM EDTA, 10 mM Na_4_P_2_O_7_, 10% Glycerol). The proteins on the FLAG beads were eluted with 30 μl of 1 mg/ml FLAG peptide and were ready for *in vitro* MT binding and gliding assays.

### Immunoblotting

Immunoblotting experiments were performed as described (Zhao et al., 2021). To prepare tissue lysates, freshly dissected tissues from mice were placed in ice-cold RIPA buffer (50 mM Tris-HCl, 150 mM NaCl, 1% NP-40, 0.5% Sodium Deoxycholate, 0.1% SDS, and 1 mM EDTA, pH7.5) supplemented with protein inhibitors cocktail. The tissues were homogenized using the homogenizer Precellys 24 system (Bertin, France). The lysates were cleared by centrifugation at 13,200 rpm at 4°C for 15 min, and mixed with an equal volume of 2 × SDS-PAGE loading buffer (100 mM Tris-HCl, 4% SDS, 20% glycerol, 2% 2-mercaptoethanol, 0.1% bromophenol blue, pH 6.8). For cultured cells, cells were washed with pre-warmed PBS and lysed in 2 × SDS-PAGE loading buffer. The lysates were boiled at 100°C for 10 min and then subjected to immunoblotting.

### *In-vitro* MT binding and gliding assays

To assemble the imaging chamber, two pieces of double-slided tape were adhered to a clean microscope slide and a silanized 18 mm × 18 mm glass coverslip was placed and pressed onto the double-sided tape, creating a chamer between the coverslip and the microscope slide as previously described (Stanhope and Ross, 2015). For MT binding assay, a solution containing 10 μg/μl of Streptavidin was diluted in 50 ul of reaction buffer and pipetted into the imaging chamber, followed by immobilization for 20 min and blocking with 1% (w/v in ddH_2_O) Pluronic F127 for 10 min. For MT gliding assay, the purified kinesin proteins were nonspecific adsorbed onto the surface of a coverslip and immobilized.

To perform MT binding assay, rhodamine-labeled MTs were polymerized using a mixture containing 5% rhodamine-labeled tubulin (Cytoskeleton, TL590M), 5% biotin-labeled tubulin (Cytoskeleton, T333P), and 90% unlabeled tubulin (Cytoskeleton, T240) in BRB80 buffer (80 mM PIPES, 1 mM EGTA,1 mM MgCl_2_, pH 6.8) supplemented with 1 mM GTP. The mixture was then incubated at 37℃ for 30 min. Next, 10 μM taxol was added, and the incubation continued at 37°C for an additional 30 min to stabilize the MTs. The sample was subsequently centrifuged at 13, 200 rpm for 25 min to remove free tubulin. The polymerized MTs were diluted in BRB80 buffer and introduced into an imaging chamber, allowing them to adsorb to the coverslips via streptavidin. Subsequently, the motor proteins, along with 1 mM ATP in BRB80 buffer, were added to the polymerized MTs and imaged using a total internal reflection fluorescent (TIRF) microscope (ZEISS Cell Observer SD). To perform MT gliding assays, rhodamine-labeled GMPCPP-stabilized MT seeds were kindly donated from Prof. Bao (Institute of Biochemistry and Cell Biology, CAS) and utilized as previously described (Diao et al., 2022; Stanhope and Ross, 2015). 3 μM Kif6-CC1 or 2 μM Kif5c-MD in BRB80 supplemented with 1 mM ATP and incubated for 5 min to pre-coated the glass surfaces. Subsequently, rhodamine-labeled GMPCPP-stabilized MT seeds plus 1 mM ATP in BRB80 buffer were introduced to examine the gliding ability of the MTs along the coverslip surface. The dynamic gliding of MTs was imaged using a TIRF microscopy at 0.72-s interval for per frame.

### Transmission electron microscopy

The tissues were fixed in 2.5% glutaraldehyde/PBS at 4°C overnight, washed with PBS, and treated with 1% OsO_4_ for 1.5 h at r.t.. The tissues were dehydrated with a graded series of ethanol and further dehydrated with acetone. Subsequently, the samples were embedded in Epon 812 resin at 60°C for 48 h. Thin sections of 70 nm thickness were obtained using an ultramicrotome (Leica EM PACT2) and stained with 2% uranyl acetate for 15 min, followed by 1% lead citrate for 5 min. The images were captured using a FEI Tecnai G2 Spirit Twin transmission electron microscope and analyzed using Fiji software.

### Coronal brain section

Coronal brain slices with a thickness of 250 μm were achieved using a vibratome sectioning technique, as previously described (Chen et al., 2020). Mice were deeply anesthetized with 360 mg/kg body weight of avertin administered via intraperitoneal injections. Transcardial perfusion was performed using PBS followed by 4% PFA in PBS, utilizing an injection pump (Smiths medical, WZS-50F6, 250 ml/h). Whole brains were dissected from the mice and cut using a vibratome to obtain coronal slices with a thickness of 250 μm. These slices were placed into 35-mm glass-bottom dishes (Cellvis, D35-20-1.5-N) and were imaged with an Olympus SZX16 Stereo Microscope.

### Reproductive performance test

Mating test was perform as previously described (Miyata et al., 2020). 2-3 month-old *Kif6*^-/-^ and *Kif9*^-/-^ male mice with normal appearance were individually caged with two age-appropriate heterozygous female mice for four months and two *Kif6*^-/-^ and *Kif9*^-/-^ female mice were separately caged with one heterozygous male mice, and then were checked regularly every several days. The number of pups was counted during the period. Three independent experiments were carried out.

### Phylogenomic analysis and multiple alignment

The phylogenetic tree and taxonomic groups were constructed based on literatures and Taxonomy Browser of National Center for Biotechnology Information (NCBI) (Cetkovic et al., 2018; Mukherjee and Brocchieri, 2013). The protein database of NCBI were searched to identify genes with orthologous proteins of Kif6 and Kif9 across a wide range of species (such as mammals, birds, amphibians, fishes, jawless vertebrates, cephalochordates, urochordates, echinoderms, nematodes, mollusks, flatworms, cnidarians, placozoans and protozoans) using the full-length murine protein sequences of Kif6 and Kif9 as queries.

The selected sequences were aligned using COBALT searches with default settings. Genebank accession numbers of Kif6 and Kif9 orthologues were provided in Supplemenatry Figure 1a. Only alignment columns without gaps were colored, with highly conserved positions shown in red and less conserved positions shown in blue.

### Analysis of BB rotational polarity

BB rotational polarity was analyzed as previously described (Guirao et al., 2010; Mirzadeh et al., 2010; Ryu et al., 2021). In TEM images, the BB rotational polarity was determined by drawing a unit vector from the center of a BB to the vertex of its BF. The mean vector length within a given field in each full-size electron micrograph (2,048 pixels × 2,048 pixels) was calculated using Microsoft Excel software (Microsoft). In 3D-SIM images, the BB rotational polarity was determined by drawing a unit vector from the geographic center of Cep164 ring (BB distal end marker) to the center of either Centriolin or γ−tubulin signals (BF marker). The mean vector length within a cell was calculated using Excel software (Microsoft) to demostrate the extent of the rotational polarity.

### Quantification and statistical analysis

BB numbers were assessed without knowledge of the genotypes and were measured from 3D-SIM images using Cep164-positive ring as a BB marker. The BB center and cell border center were determined using the default setting of the “ROI Manager” function in Fiji.

All experiments, unless otherwise stated, were independently repeated at least twice for both microscopic or biochemical results. Statistical results are presented as mean ± SD. Unpaired two-tailed student’s *t*-test was performed using GraphPad Prism 9.0 software, and differences were considered significant at *P* < 0.05. Data analyses were performed in a blinded manner, with the researchers performing the quantification being unaware of the genotype.

## Acknowledgements.

The authors thank Profs. Lan Bao and Dr. Lei Diao (Shanghai Institute of Biochemistry and Cell Biology, CAS) for the reagents and technical assistance, Profs. Xin Liang (Tsinghua University) and Profs. Wei Feng (Institute of Biophysics, CAS) for technical help and valuable discussions, and the staff members of the Integrated Laser Microscopy System at the National Facility for Protein Science in Shanghai (NFPS) (Shanghai Advanced Research Institute, CAS) and core facilities of the Center for Excellence in Molecular Cell Science (CEMCS) for instrumental and technical supports. This work was supported by National Natural Science Foundation of China (31991192, 32230027 and 32270725).

## Declaration of interests

The authors declare no competing financial interests.

## Author contributions

X.Y., X.Z., and Dong L. conceived and supervised the project; C.F., X.P. and Di L. performed the research; Dong L. provided GI-SIM; Y.C., L.L. and Q.G. assisted with experiments; X.Y., X.Z., C.F. and X.P. collectively analyzed data and wrote the manuscript, with input from all authors.

## Legends for Supplementary Figures

**Supplementary Figure 1.**
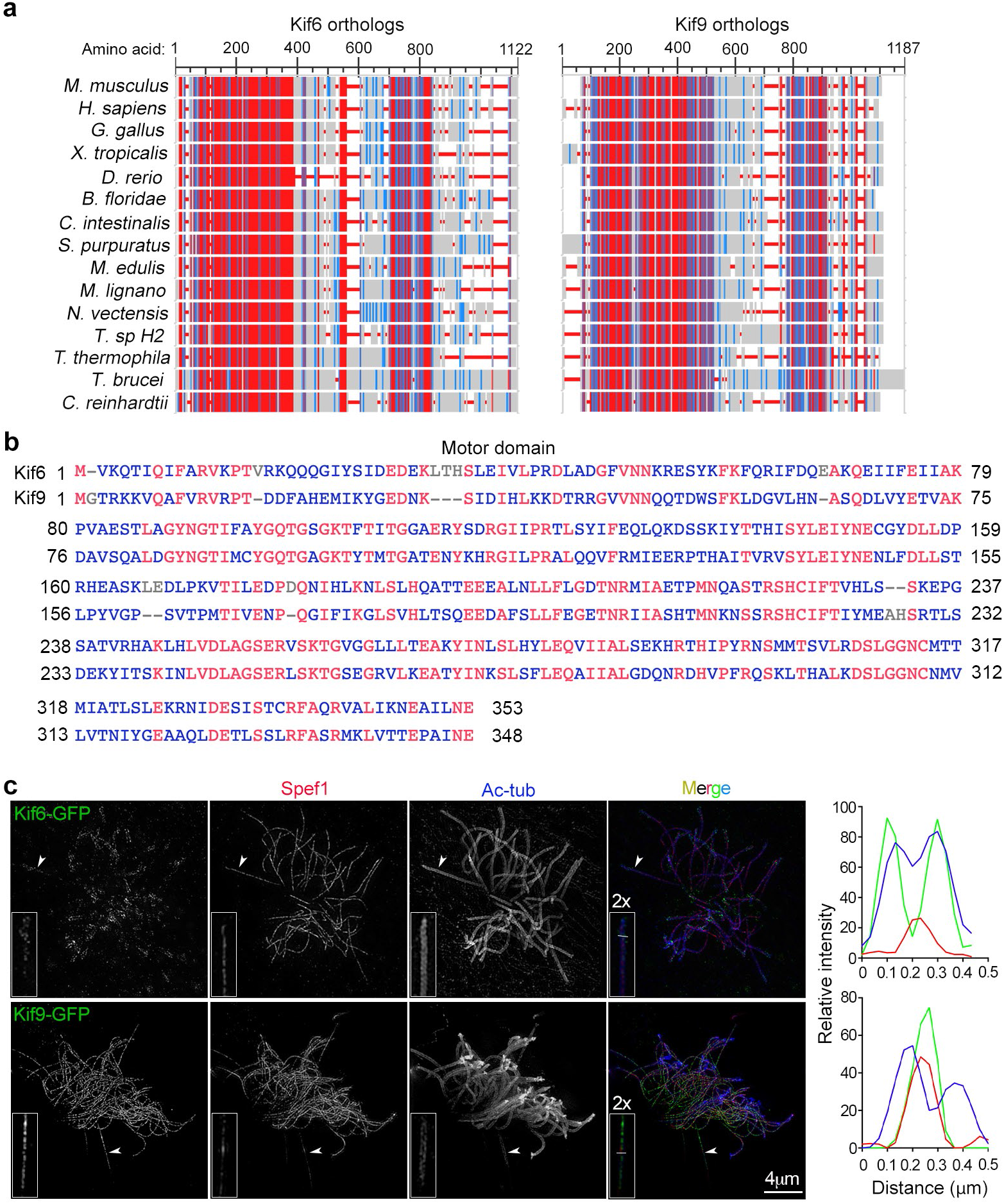
Conservations and ciliary localizations of Kif6 and Kif9 (related to Fig. 1). **(a)** Similarities of Kif6 (left) and Kif9 (right) orthologues, obtained using constraint-based multiple alignment tool (COBALT) in NCBI. Highly conserved positions are shown in red and less conserved positions shown in blue. Protein sequences of Kif6 orthologues used were: mouse (NP_796026.2), human (NP_659464.3), chicken (XP_003640993.5), frog (NP_001106482.1), zebrafish (NP_001070899.1), lancelet (XP_035685117.1), ascidian (XP_002124970.1), sea urchin (XP_030854411.1), mussel (CAG2221785.1), flatworm (PAA61917.1), sea anemone (XP_032228212.2), *Trichoplax* (RDD42533.1), *Tetrahymena* (XP_001024804.2), *Trypanosoma* (XP_846346.1) and green algae (XP_042928646.1). Protein sequences of Kif9 orthologues used were: mouse (NP_001157041.1), human (NP_001400904.1), chicken (XP_040522129.1), frog (XP_012821147.1), zebrafish (XP_001922460.1), lancelet (XP_035663761.1), ascidian (XP_018668035.1), sea urchin (XP_030830023.1), mussel (CAG2242108.1), flatworm (PAA61899.1), sea anemone (XP_001636554.1), *Trichoplax* (RDD45058.1), *Tetrahymena* (XP_001022313.1), *Trypanosoma* (RHW71316.1) and green algae (P46870.1). **(b)** Sequence alignment of Kif6 and Kif9 motor domains, generated by the COBALT sequence alignment program using default parameters. GenBank accession numbers of proteins were as follows: NP_796026.2 (Kif6) and NP_001157041.1 (Kif9). The motor domains were identified using SMART. **(c)** Localizations of exogenous Kif6 and Kif9 in motile cilia. Cultured mEPCs were treated as indicated in Figure 1g to express Kif6-GFP or Kif9-GFP and imaged by 3D-SIM. Spef1 and acetylated tubulin (Ac-tub) served as markers for the CP and the axoneme, respectively. Typical cilia (arrowheads) were magnified by 200% to highlight details. Line scans were generated at the indicated positions by white lines in merged images.

**Supplementary Figure 2.**
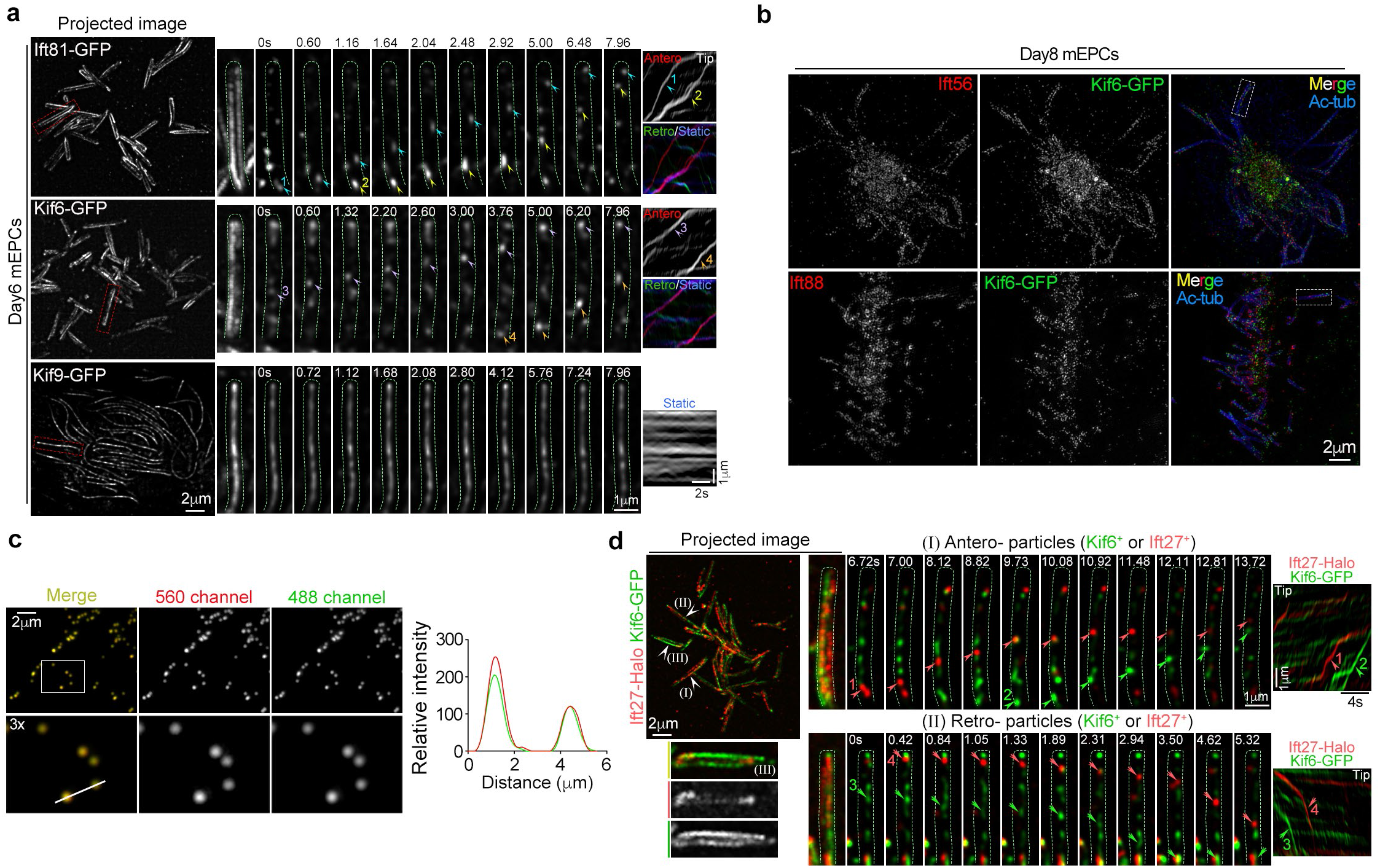
Kif6 moves along the axoneme with or without IFT-B particles (related to Fig. 2 and 3). **(a)** Kif6-GFP, but not Kif9-GFP, resembling Ift81-GFP, moved along the axonemal DMTs. The trafficking trajectories of GFP-tagged puncta were visualized through projections of the first 200 frames. Typical cilia framed by dashed red lines in projected images were magnified by 300% to showed details. Dashed green lines were used to outline the cilia. The clear and traceable GFP-tagged particles were marked and corresponding kymographs were presented. **(b)** Co-immunostainning Kif6-GFP with IFT proteins (Ift56 and Ift88) imaged by 3D-SIM. The framed cilia were magnified and presented in Figure 3a. Ac-tub marked ciliary axoneme. (**c**) Calibration of the microscope. Instrument calibration was conducted routinely with fluorescent beads before dual-color GI-SIM imaging experiments. A line scan was performed along the white line to determine the alignment. **(d)** Ciliary trafficking trajectories of GFP/Halo-tagged puncta were projected by the first 200 frames. Typical cilia pointed were magnified by 300% to show details. Dashed green lines were used to outline the cilia. Kif6^+^ particles or Ift27^+^ particles are indicated by colored arrowheads or arrows to respectively show anterograde or retrograde transports from image time sequences. Corresponding kymographs were presented.

**Supplementary Figure 3.**
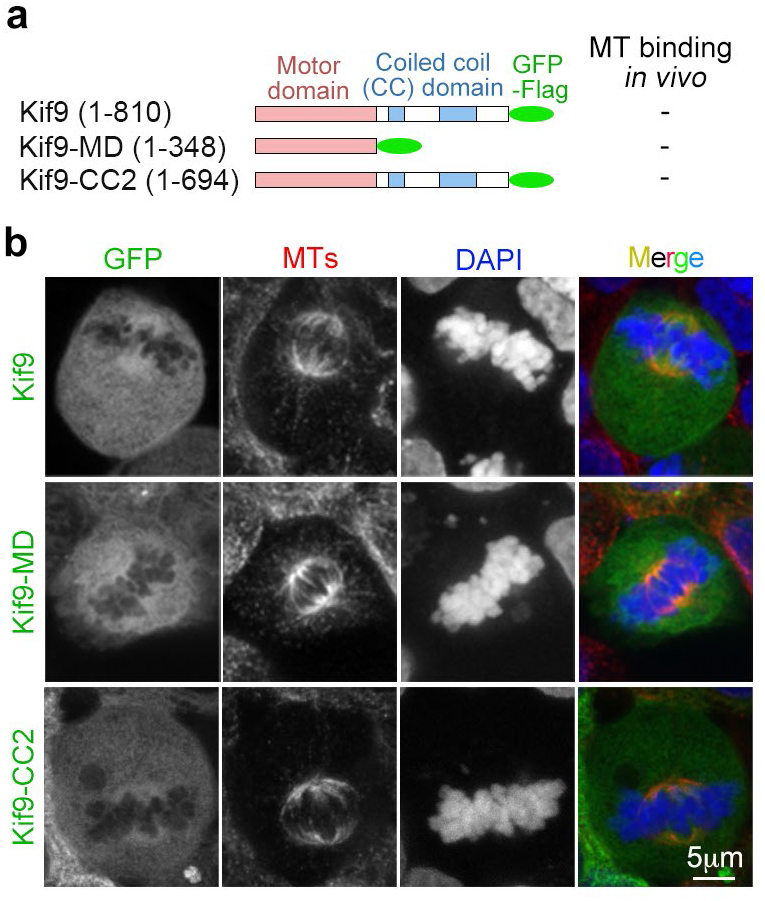
Kif9 does not exhibit MT associations in HEK293T cells (related to Fig. 4). **(a)** Diagrams of Kif9 and its deletion constructs showing their MT-binding ability. **(b)** Kif9 and its deletion constructs displayed no spindle MTs binding. MTs were visualized by an anti-α-tubulin antibody. Chromosomes and nuclei were stained with DAPI, a DNA-specific dye.

**Supplementary Figure 4.**
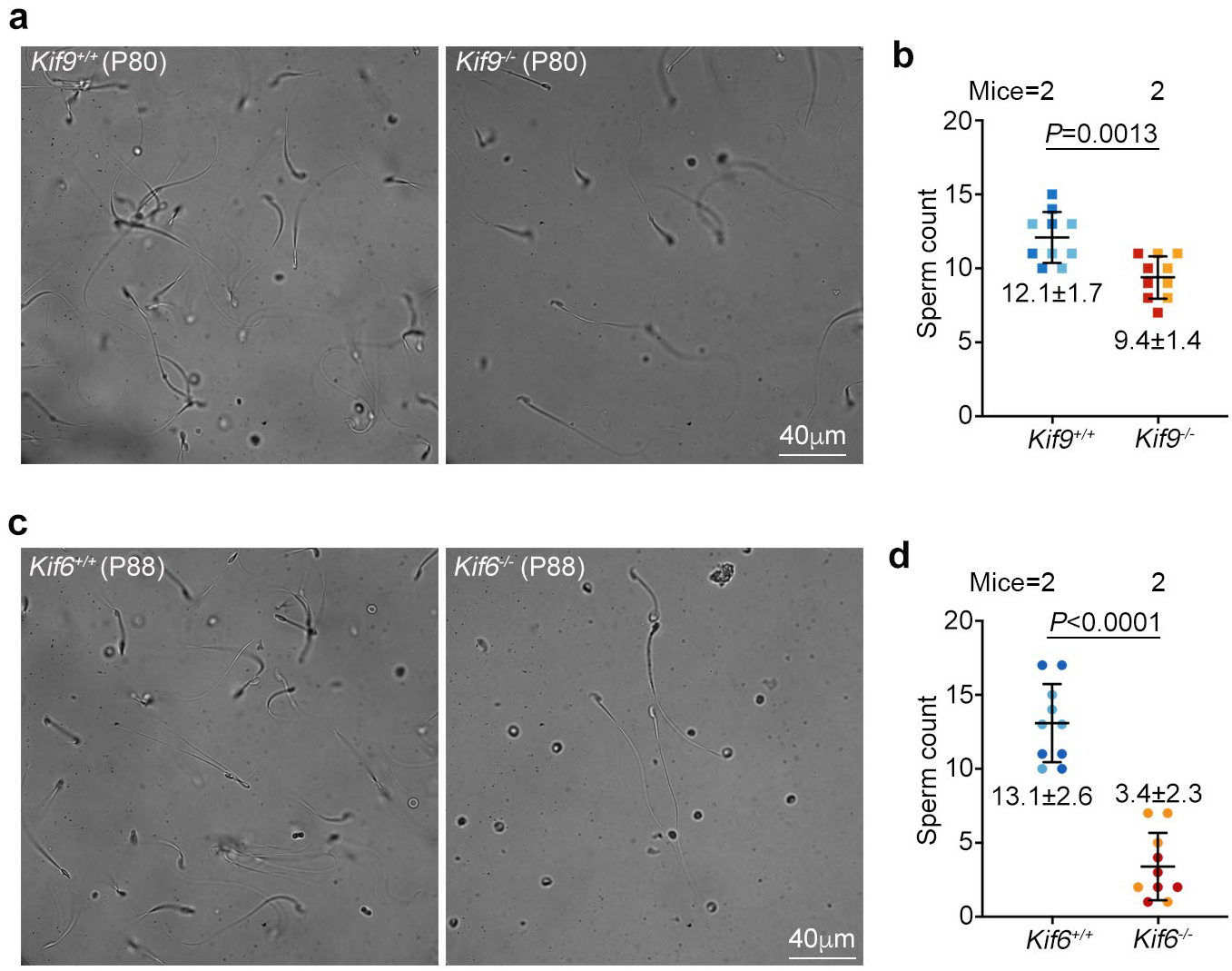
*Kif6-*deficient and *Kif9-*deficient males show decreased sperm number (related to Fig. 5). Sperm were released from cauda epididymis, followed by a spinning disk confocal microscope at a fixed z-plane. Representative images (**a, c**) were from cauda epididymis of littermates. Quantification results (**b, d**) from two mice each genotype (5 full-size micrographs per mouse) were pooled together (mean ± SD plus sample dots) and subjected to unpaired two-tailed student’s *t*-test.

**Supplementary Figure 5.**
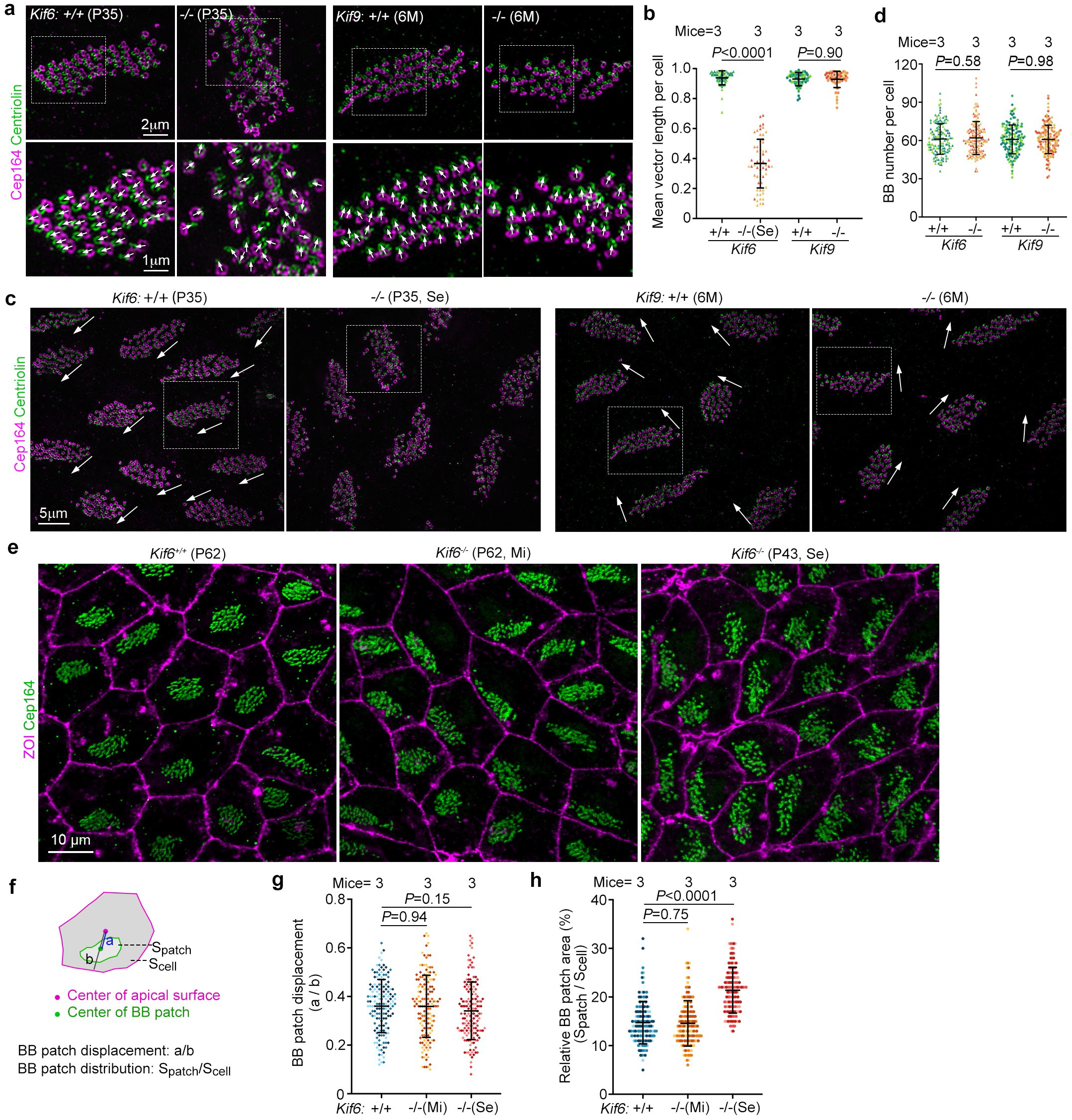
*Kif6* deficiency disrupts rotational polarity but not the translational polarity of BBs in ependyma (related to Fig. 7). Se: Severe hydrocephalus mice with obvious dome-shaped skull. Mi: mild hydrocephalus mice with normal morphological appearance but enlarged ventricles. **(a-c)** BBs in *Kif6^-/-^* but not in *Kif9^-/-^*ependymal cells displayed impaired rotational polarity. Ependymal tissues from the indicated littermates were co-immunostained for Cep164 to visualize BBs and Centriolin to visualize BFs. Framed regions in (**a**) were maginified to show BFs orientations (arrows). Framed regions in (**c**) were presented in (**a**). Arrows in (**c**) indicate rotational polarities of the wild-type and *Kif9^-/-^* BB patches, which broadly point to one direction in the microscopic field. Quantification of the rotational polarity **(b)**. Results from three mice per genotype were presented as mean ± SD plus sample dots and subjected to unpaired two-tailed student’s *t*-test. Mice used were: two 6-month-old (6M) and one 7-month-old (7M) mice for *Kif9^-/-^*and three of their wild-type littermates; three P35 mice with severe hydrocephalus for *Kif6^-/-^* and three of their wild-type littermates. 20 ependymal cells were scored for each mouse. **(d)** BB number in ependymal cells was not altered by *Kif6* or *Kif9*-deficiency. 50 cells were measured per mouse. (e-h) *Kif6* deficiency did not impair the transitional polarity of BBs. Ependymal tissues from two P62 and one P68 *Kif6^-/-^* mice of mild hydrocephalus, three severe hydrocephalus (P34, P38 and P43) and three wild-type mice (P62, P38, and P43) were immunostained for ZO1 and Cep164 to visualize cell borders and BBs, respectively (**e**). BB patch displacement relative to the cellular geographic center (**g**) and relative BB patch area (**h**) were quantified as illustrated in (**f**). 50 ependymal cells were scored for each mouse.

## Legends for supplementary movies

**Movie 1. Bidirectional motility of ciliary Ift81-GFP puncta (related to Fig. 2c).** mEPCs were infected with adenovirus at day-1 and imaged during day 5-8 by GI-SIM. Images were recorded at 25 frames per sec (fps) and the playback speed is 20 fps. Typical cilia framed were magnified by 300% to show details of Ift81-GFP puncta. The clear particles were marked by arrowheads or arrows to show anterograde or retrograde transports, respectively.

**Movie 2. Bidirectional motility of ciliary Kif6-GFP puncta (related to Fig. 2d).** mEPCs were treated, imaged, and presented as described in Movie 1. Typical cilia framed were magnified by 300% to show detailed bidirectional transports of Kif6-GFP puncta. The traceable particles were indicated by arrowheads or arrows to show anterograde or retrograde transports, respectively.

**Movie 3. Kif9-GFP does not exhibit progressive movement along the cilia (related to Fig. 2e).** mEPCs were treated, imaged, and presented as described in Movie 1. Typical cilia framed were magnified by 300% to show details.

**Movie 4. Ciliary Kif6-GFP puncta move together with IFT-B particles (related to Fig. 3b).** mEPCs were infected with adenovirus expressing Kif6-GFP and Ift27-Halo, and dual-color time-lapse live imaging was taken by GI-SIM. Images were recorded at 14 fps and the playback speed is 20 fps. Typical cilia pointed were magnified by 300% to display clear anterograde (arrowheads) and retrograde (arrows) transports of Ift27^+^Kif6^+^ duo-particles. mEPCs were infected with adenovirus at day-1 and imaged during day 5-8 by GI-SIM. Images were recorded at 25 fps and the playback speed is 20 fps.

**Movie 5. Ciliary Kif6-GFP puncta move without IFT-B particles (related to Fig. 3c).** mEPCs were treated, imaged, and presented as described in Movie 4. Typical cilia pointed were magnified by 300% to display traceable anterograde (arrowheads) and retrograde (arrows) transports of Kif6^+^ particles or Ift27^+^ particles (related to Fig. 3c).

**Movie 6. The MT gliding of Kif6-CC1 (related to Fig. 4h).** Kif5c-MD served as a positive control. 2 μM Kif6-CC1 (left) or 2 μM Kif5c-MD (right) respectively in BRB80 supplemented with 1 mM ATP. Yellow arrowheads indicate the gliding MTs. The assay was acquired at 0.72s/frame. The playback speed is 10 fps.

**Movie 7. A representative sperm motility from *Kif9^+/+^* and *Kif9^-/-^* male mice.** Sperms were released from cauda epididymis, followed by live imaging with a spinning disk confocal microscope at a fixed z-plane and at 25-ms intervals. The playback speed is 20 fps. One pair of P80 wild-type (left) mice and *Kif9*^-/-^ (right) litermates were presented.

**Movie 8. A representative sperm motility from *Kif6^+/+^* and *Kif6^-/-^* male mice.** Sperm were released, imaged, and presented as described in Movie 7. One pair of P88 wild-type (left) mice and *Kif6*^-/-^ (right) litermates were presented.

**Movie 9. Representative ciliary beat patterns in *Kif6^+/+^*ependyma (related to Fig. 6f).** The P31 *Kif6^+/+^* ependyma was freshly dissected and labeled with SiR-tubulin, followed by live imaging with a spinning disk confocal microscope at a fixed z-plane and at 10-ms intervals. The playback speed is 5 fps. Images of the first three frames are presented in Figure 6f.

**Movie 10. Representative ciliary beat patterns in *Kif6^-/-^* ependyma (related to Fig. 6f).** The P31 *Kif6^-/-^* ependyma was treated, imaged, and presented as described in Movie 9. The mouse used was a littermate of the one in Movie 9, and exhibited severe hydrocephalus. Images of the first three frames are presented in Figure 6f.

**Movie 11. Representative liquid flows were driven by ciliary beat in *Kif6^+/+^* ependyma (related to Fig. 6i).** Fresh ependyma from a P36 wild-type mouse were incubated with SiR-tubulin to label cilia and fluorescent latex beads as flow indicators. Beads motilities were imaged with a spinning disk confocal microscope at 100-ms intervals and at a fixed z-plane. The playback speed is 5 fps. Trajectories of 8 traceable, rapidly-moving aggregated beads in the first 500 ms were shown. The projections of bead movements are presented in Figure 6i.

**Movie 12. Representative liquid flows were driven by ciliary beat in *Kif6^-/-^* ependyma (related to Fig. 6i).** Fresh ependymal tissues from a P36 *Kif6^-/-^* mouse with severe hydrocephalus were treated, imaged, and presented as described in Movie 13. Trajectories of 13 traceable, rapidly-moving aggregated beads in the first 500 ms were shown. The projections of bead movements are presented in Figure 6i.

## Legends for Supplementary Tables

**Supplementary Table 1 Sequences of primers and gRNAs.**

**Supplementary Table 2 List of antibodies.**

## Notes

### Competing Interest Statement

The authors have declared no competing interest.

### Summary of Updates

Figure 4 revised.

